# Pathogenic tau inhibits synaptic plasticity by blocking eIF4B-mediated local protein synthesis

**DOI:** 10.1101/2025.09.11.675671

**Authors:** Grant Kauwe, Doyle Lokitiyakul, Ivy L. Wong, Kevin Schneider, Varun Sridhar, Durai Sellegounder, Yani Y. Ngwala, Lei Yao, Jackson H. Chen, Kristeen A. Pareja-Navarro, Alissa L. Nana, Salvatore Spina, William W. Seeley, Lea T. Grinberg, Eric Verdin, David Furman, Celeste M. Karch, Li Gan, Tara E. Tracy

## Abstract

Activity-dependent modulation of synaptic strength is critical for encoding memories and it is inhibited in tauopathies including Alzheimer’s disease (AD) and Frontotemporal lobar degeneration with tau inclusions (FTLD-tau). Pathogenic tau accumulates in neurons where it obstructs synaptic plasticity. How tau blocks synaptic plasticity leading to memory loss is unclear. Here, we show that FTLD-tau inhibits plasticity by blocking activity-dependent protein synthesis in dendrites. In the plasticity-associated translatome, we identified a subset of downregulated translated mRNAs in FTLD-tau neurons that encode postsynaptic plasticity regulators. Protein synthesis was blocked by FTLD-tau binding to eIF4B which caused eIF4B dissociation from the translation initiation complex and reduced dendritic eIF4B levels. Inhibiting the tau-eIF4B interaction or enhancing eIF4B levels in FTLD-tau neurons restored local protein synthesis and synaptic plasticity. Together, this suggests that pathogenic tau binding to eIF4B disables the local synthesis of plasticity-related proteins that drive synapse strengthening and memory formation.

## Introduction

As highly dynamic structures, synapses modify their strength in response to neuronal activity and their unique molecular properties are essential for plasticity in the brain. A persistent strengthening of glutamatergic synapses during plasticity, termed NMDA receptor (NMDAR)-dependent long-term potentiation (LTP), is a fundamental mechanism in the hippocampus that underlies the formation of new memories^1^. Impaired LTP in hippocampal neurons is a common pathophysiology that coincides with memory decline across tauopathy mouse models, including Alzheimer’s disease (AD) mouse models^2–10^. Tauopathies are age-related neurodegenerative diseases that are characterized by the aggregation of pathological tau protein in the brain which correlates with cognitive decline. In these diseases, the accumulation of toxic tau can drive pathogenesis in neurons through progressive and complex mechanisms^11^. Mutations in the *MAPT* gene that encodes tau cause Frontotemporal lobar degeneration with tau inclusions (FTLD-tau)^12^, and pathogenic forms of tau associated with tauopathy include oligomeric tau^13^ and tau with abnormal post-translational modifications^14^. These forms of pathogenic tau inhibit LTP at synapses and cause tauopathy-related memory deficits in mouse models^2–4,15–20^. Notably, LTP impairment occurs early in the progression of tau-mediated pathogenesis preceding the later onset of neurodegeneration^2,3^.

In addition to its role as a microtubule binding protein, growing evidence suggests that tau interacts with broader protein networks in neurons which may contribute to neuronal pathophysiology and deterioration in disease^11,21–23^. Thus, pathogenic tau can affect neurons and their function via a range of mechanisms at different stages of disease progression. This poses a challenge in narrowing down the specific role of tau in causing the synapse pathophysiology that drives memory loss in tauopathy. The series of postsynaptic molecular signaling events and processes that are required for the expression of LTP are also complex and multifaceted^24^. Consequently, the mechanism by which pathogenic tau inhibits LTP at synapses has not been determined.

Protein synthesis is required for LTP and memory, and it involves the activity-dependent increase of mRNA translation in dendrites^25–31^. The localized synthesis of proteins in dendrites plays a critical role in the enhancement of postsynaptic strength for LTP^32–34^. Here, we report that FTLD-tau blocks activity-dependent protein synthesis in dendrites which is required for LTP expression at synapses. With RNA sequencing (RNA-seq) of polysome-bound mRNAs, we probed the LTP-associated translatome in human neurons and identified the subset of synapse-related transcripts that were dysregulated in FTLD-tau neurons. Finally, we uncovered FTLD-tau binding to eIF4B as a key mechanism that inhibits protein synthesis during plasticity by hindering eIF4B’s function in activity-dependent translation initiation.

## Results

### Dysregulated LTP expression in FTLD-tau human neurons

To investigate synaptic plasticity in human neuron models, we adapted a chemical method of LTP induction (cLTP) using glycine in the absence of magnesium to activate synaptic NMDARs which leads to increased postsynaptic AMPA receptor (AMPAR) insertion and strengthened synaptic transmission^35^. Human glutamatergic neurons with functional synapses were assessed 6-8 weeks after the differentiation of induced pluripotent stem cells (iPSCs) with the genetic integration of an inducible neurogenin-2 (NGN2) transgene^36^. Surface immunolabeling of GluA1-containing AMPARs was performed on human neurons after cLTP induction to monitor AMPAR trafficking to the postsynaptic surface at synapses marked by co-localization with presynaptic vGluT1. Levels of AMPARs were significantly increased at the postsynaptic surface in control human neurons with wild-type tau (tauWT) 30 min after cLTP compared to unstimulated neurons (Fig. 1a,b). The trafficking of AMPARs after cLTP induction was blocked by the NMDAR antagonist APV (Extended Data Fig. 1a,b), which confirmed the NMDAR-dependence of LTP expression in human neurons. We next tested AMPAR trafficking during LTP in isogenic neurons carrying the V337M FTLD-causing homozygous *MAPT* mutation (tauV337M_iso_) introduced by CRISPR editing^37^. In contrast to tauWT neurons, surface immunolabeling of AMPARs at the synapses of tauV337M_iso_ neurons was not significantly changed following cLTP compared to unstimulated neurons (Fig. 1a,b), suggesting that LTP-induced postsynaptic AMPAR trafficking was inhibited in tauV337M_iso_ neurons. To probe for changes in synaptic NMDARs that could affect LTP, we analyzed immunolabeling of the GluN1 NMDAR subunit at synapses but found that GluN1-containing NMDARs at synapses were not altered in tauV337M_iso_ neurons compared to tauWT neurons (Extended Data Fig. 1c,d). The density of synapses was not significantly different between tauV337M_iso_ and tauWT neurons (Fig. 1c,d). Likewise, both cultures had similar MAP2-immunolabeled dendrite coverage, postsynaptic PSD-95 levels and presynaptic Synapsin levels (Extended Data Fig. 1e-g), indicating that the overall synaptic and dendritic structures of tauV337M_iso_ neurons were intact. We further tested LTP in human neurons derived from FTLD-tau patient iPSCs carrying a heterozygous V337M mutation (tauV337M_pat_) and isogenic CRISPR edited iPSCs with the tauV337M mutation corrected (tauV337V_iso_)^38^. Postsynaptic AMPAR recruitment during LTP was blocked in tauV337M_pat_ neurons, but it was restored in the corrected tauV337V_iso_ neurons which had significantly increased levels of AMPARs at the postsynaptic surface after cLTP induction compared to both tauV337M_pat_ neurons and to unstimulated tauV337V_iso_ neurons (Fig. 1e,f). Levels of vGluT1, a presynaptic vesicular protein, were not altered by cLTP or by the V337M mutation (Extended Data Fig. 1h,i).

**Fig. 1.**
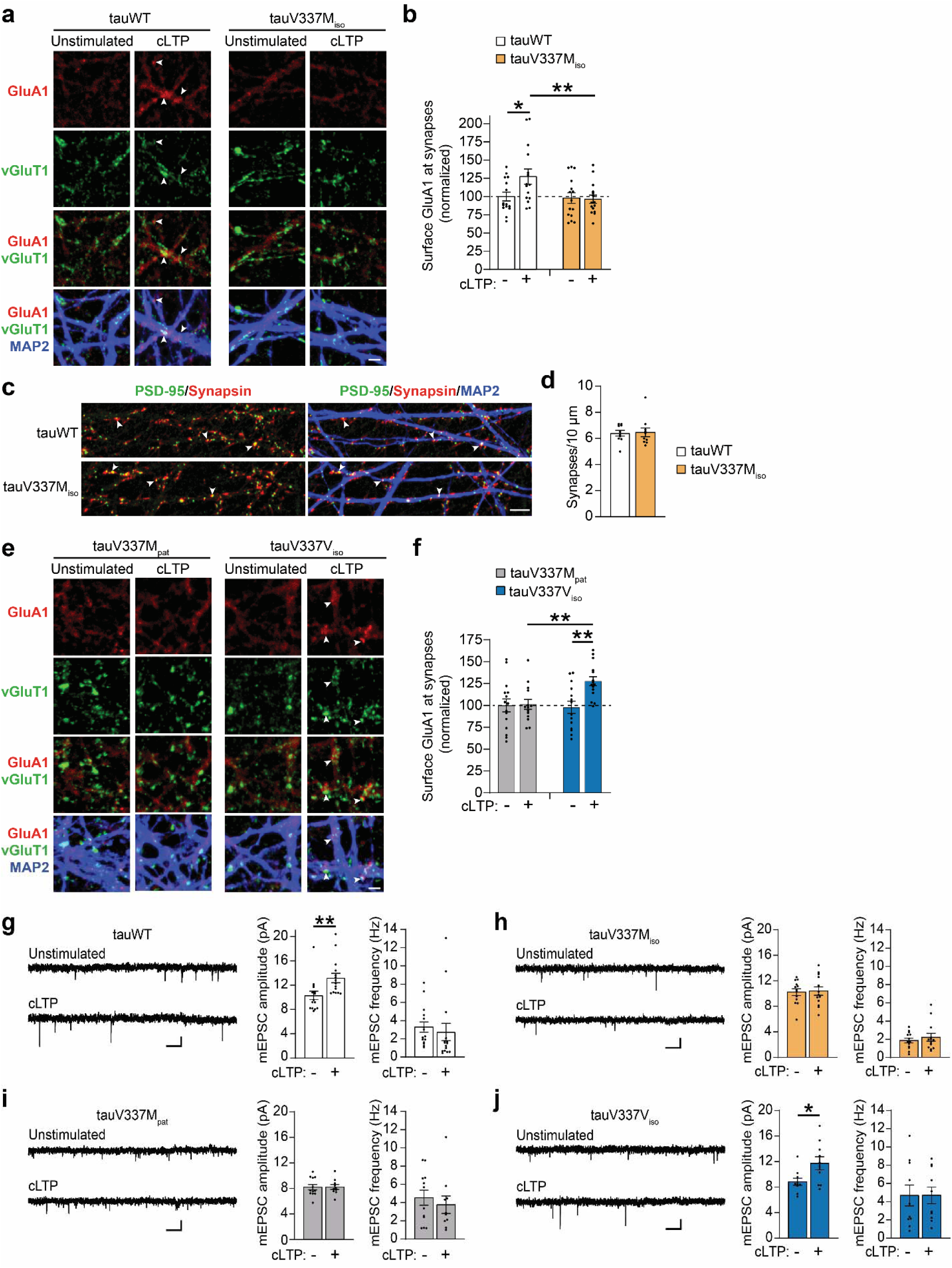
FTLD-tau blocks the recruitment of postsynaptic AMPARs required for LTP expression in human neurons. (**a**) Immunolabeling of surface GluA1-containing AMPARs (red) that co-localized with vGluT1 puncta (green) at synapses on MAP2-labeled dendrites (blue) of human neurons. Neurons were either unstimulated or treated for chemical long-term potentiation (cLTP) induction. Synapses on tauWT neurons, but not isogenic tauV337M_iso_ neurons, had increased levels of surface GluA1-containing AMPARs (white arrowheads) at 30 min after cLTP induction. Scale bar, 2 μm. (**b**) Quantification of surface GluA1 immunolabeling intensity that overlapped with the vGluT1 puncta along the dendrites of tauWT and tauV337M_iso_ neurons (n = 15 images/group; * p < 0.05, ** p < 0.01, two-way ANOVA, Bonferroni post-hoc analyses). Values were normalized to the mean GluA1 intensity at synapses on unstimulated tauWT neurons. (**c**) Confocal images show co-localized immunolabeling of postsynaptic PSD-95 (green) and presynaptic Synapsin (red) at synaptic connections (yellow, arrowheads) on human neuron dendrites marked by MAP2 (blue). Scale bar, 5 μm. (**d**) Quantification of the density of co-localized PSD-95 and Synapsin puncta marking synapses on dendrites (n = 10 images/group; not significant, Student’s t-test). (**e**) Images show immunolabeling of surface GluA1-containing AMPARs (red) co-localized with vGluT1 puncta (green) on MAP2-labeled dendrites (blue) in human neurons from a tauopathy patient (tauV337M_pat_) and isogenic neurons with the V337M mutation on tau corrected (tauV337V_iso_). Synapses on tauV337V_iso_ neurons demonstrated cLTP-induced GluA1 postsynaptic insertion (arrowheads), but AMPAR trafficking was inhibited in tauV337M_pat_ neurons. Scale bar, 2 μm. (**f**) Quantification of surface GluA1 immunolabeling intensity that overlapped with vGluT1 puncta along dendrites in tauV337M_pat_ and tauV337V_iso_ neurons (n = 14-15 images/group; ** p < 0.01, two-way ANOVA, Bonferroni post-hoc analyses). Values were normalized to the mean GluA1 intensity at synapses on unstimulated tauV337M_pat_ neurons. (**g-j**) Representative traces and analyses of miniature excitatory postsynaptic currents (mEPSCs) recorded from individual neuron cultures that were unstimulated or that received cLTP treatment. The mean mEPSC amplitude and frequency was quantified from each culture with and without LTP induction. The increased mEPSC amplitude following cLTP in (**g**) tauWT (n = 15 neurons/group; ** p < 0.01, Student’s *t*-test) and (**j**) tauV337V_iso_ neurons (n = 10 neurons/group; * p < 0.05, Student’s *t*-test) was blocked in both (**h**) tauV337M_iso_ (n = 13 neurons/group; not significant, Student’s *t*-test) and (**i**) tauV337M_pat_ neurons (n = 10-12 neurons/group; not significant, Student’s *t*-test). Scale bars: 10 pA and 250ms. Values are given as means ± SEM.

To measure synapse strength directly, whole-cell patch-clamp recordings of miniature excitatory postsynaptic currents (mEPSCs) were performed on tauWT neurons. The increase in mEPSC amplitude after LTP induction reflects the potentiation of synaptic strength in neuronal cell culture models^35,39^. There was a significant increase in mEPSC amplitude in tauWT neurons more than 30 min after cLTP treatment compared to unstimulated neurons with no significant effect on mEPSC frequency (Fig. 1g), which is consistent with the potentiation of synaptic transmission through postsynaptic AMPAR insertion. The mEPSCs recorded from tauV337M_iso_ and tauV337M_pat_ neurons exhibited comparable amplitudes and frequencies whether the neurons were unstimulated or cLTP-treated (Fig. 1h,i), indicating impaired LTP. However, tauV337V_iso_ neurons maintained significantly increased mEPSC amplitude after cLTP induction (Fig. 1j). Thus, the V337M FTLD-tau mutation inhibits the postsynaptic expression of LTP in human neurons.

### Pathogenic tau blocks activity-dependent local protein synthesis

Local protein synthesis in dendrites is required for memory and LTP expression at synapses^33,40,41^. We next investigated the role of local protein synthesis in tau-mediated LTP impairment. First, we probed whether translation machinery was present in the dendrites of human neurons. RPS6, a component of the 40s ribosomal subunit, and the 7-methylguanosine cap (m^7^G-Cap), which is added to mRNAs for cap-dependent translation, were co-localized with MAP2 immunolabeling in dendrites of tauWT neurons (Extended Data Fig. 2a,b), supporting the presence of cap-dependent translation initiation factors in dendrites. Consistent with the detection of polysomes in dendrites of hippocampal neurons^42^, ultrastructural visualization of synapses by electron microscopy revealed polysomes in dendrites of tauWT neurons near postsynaptic sites (Extended Data Fig. 2c). Polysomes are complexes of ribosomes that actively translate a single mRNA during either basal or activity-dependent protein synthesis^43,44^. To determine the role of *de novo* protein synthesis in LTP expression in human neurons, tauWT neurons were treated with anisomycin to block translation both during and after cLTP induction. Control vehicle-treated neurons demonstrated activity-dependent postsynaptic AMPAR insertion, but postsynaptic AMPAR trafficking was blocked in tauWT neurons treated with anisomycin (Fig. 2a,b), indicating that *de novo* protein synthesis is required for LTP expression in human neurons.

**Fig. 2.**
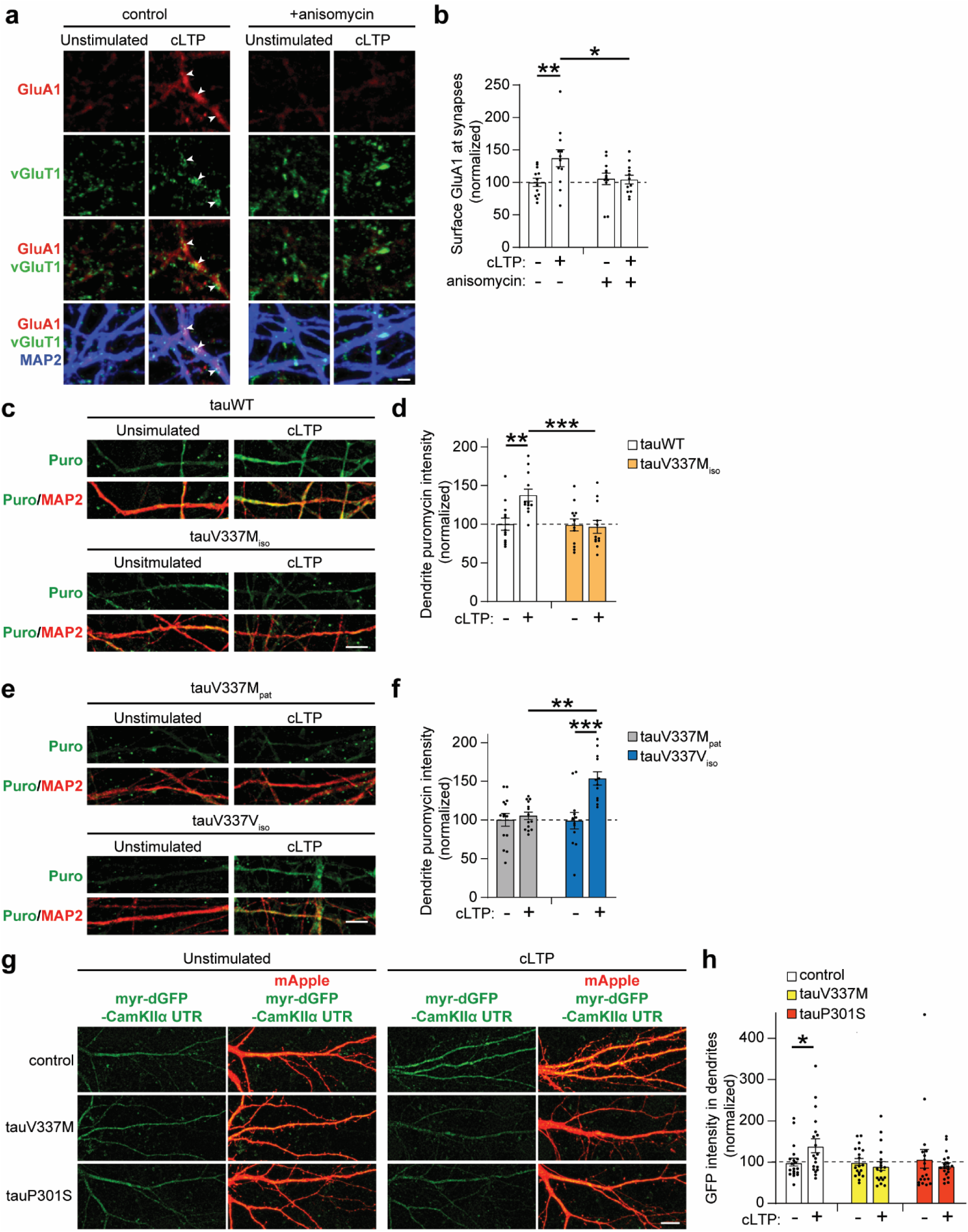
FTLD-tau blocks activity-dependent dendritic protein synthesis in human neurons. (**a**) Images of surface GluA1 (red), vGluT1 (green), and MAP2 (blue) immunolabeling in tauWT human neurons. For the duration of both unstimulated and cLTP experimental conditions, tauWT neurons were treated with either vehicle control or anisomycin (40 μm) to block protein synthesis. The postsynaptic recruitment of GluA1-containing AMPARs during LTP expression (arrowheads) was inhibited by anisomycin. Scale bar, 2 μm. (**b**) Graph of the mean surface GluA1 intensity co-localized with vGluT1 along dendrites of tauWT human neurons with and without cLTP induction and the addition of anisomycin (n = 12 images/group; * p < 0.05, ** p < 0.01, two-way ANOVA, Bonferroni post-hoc analyses). Values were normalized to the mean GluA1 intensity of the unstimulated vehicle control group. (**c**) Puromycin was applied to live neurons for 15 min to label newly synthesized proteins. Immunolabeling of puromycin (green) was detected in dendrites marked by MAP2 immunolabeling (red). The increased protein synthesis detected in tauWT neurons after cLTP was obstructed in tauV337M_iso_ neurons. Scale bar, 5 μm. (**d**) Quantification of the puromycin immunolabeling intensity measured in dendrites of tauWT and tauV337M_iso_ neurons (n = 12 images/group; ** p < 0.01, *** p < 0.001, two-way ANOVA, Bonferroni post-hoc analyses). All values were normalized to the mean puromycin intensity in dendrites of the unstimulated tauWT neurons. (**e**) Images of puromycin (green) and MAP2 (red) co-immunostaining in dendrites of tauV337M_pat_ and tauV337V_iso_ human neurons with puromycin labeling of de novo synthesized proteins. Scale bar, 5 μm. (**f**) Quantification of the puromycin immunolabeling intensity measured in dendrites of tauV337M_pat_ and tauV337V_iso_ neurons (n = 12-14 images/group; ** p < 0.01, *** p < 0.001, two-way ANOVA, Bonferroni post-hoc analyses). All values were normalized to the mean puromycin intensity in dendrites of the unstimulated tauV337M_pat_ neurons. (**g**) Images of dendrites from cultured rat hippocampal neurons co-transfected with mApple (red) and myr-dGFP-CamKIIα UTR (green), a GFP-based reporter of CamKIIα translation. Neurons with expression of human tau carrying either the V337M or P301S FTLD *MAPT* mutations and control neurons without human tau were unstimulated or treated for cLTP induction, and the GFP reporter expression in dendrites marked by mApple was monitored. Scale bar, 10 μm. (**h**) Quantification of the GFP reporter in dendrites of rat hippocampal neurons transfected with myr-dGFP-CamKIIα UTR. After cLTP induction, control neurons exhibited increased GFP levels in dendrites, but GFP levels did not change in dendrites of neurons expressing human tauV337M or tauP301S (n = 19 neurons/group; * p < 0.05, Student’s *t*-test). Analyses were normalized to the mean GFP intensity in the unstimulated control neurons. Values are given as means ± SEM.

We next investigated activity-dependent protein synthesis in the dendrites of human neurons using puromycin labeling of newly synthesized proteins. Puromycin was added to neurons 30 min after cLTP induction or to unstimulated control neurons, and newly synthesized proteins were labeled by puromycin in the neurons for 15 min. In line with reports of increased local translation of mRNAs in dendrites after LTP induction in rodent models^32,33^, we found that puromycin immunolabeling was significantly increased in the dendrites of tauWT neurons after cLTP compared to unstimulated controls (Fig. 2c,d). Interestingly, the enhancement of puromycin immunolabeling after cLTP was blocked in tauV337M_iso_ neurons (Fig. 2c,d). Similarly, tauV337V_iso_ neurons, but not tauV337M_pat_ neurons, showed a significant increase in puromycin labeling in dendrites following cLTP (Fig. 2e,f). Thus, pathogenic tau inhibits activity-dependent protein synthesis in dendrites of human neurons. Previous studies showed that basal protein synthesis was downregulated in mouse neurons with high levels of human tau pathology^45,46^. We monitored basal protein synthesis in the soma of human neurons by puromycin labeling. The intensity of puromycin labeling was not significantly different between the somas of tauWT and tauV337M_iso_ neurons and between the somas of tauV337M_pat_ and tauV337V_iso_ neurons (Extended Data Fig. 2d-g), supporting that basal somatic protein synthesis is maintained in the FTLD-tau human neurons with endogenous tau levels. Together, these results point towards a specific mechanism by which tau inhibits activity-dependent protein synthesis in dendrites that is likely distinct from mechanisms that alter basal translation associated with severe tau pathology.

LTP induction leads to the increased synthesis of Ca2+/calmodulin-dependent kinase IIα (CAMKIIα) protein in neuronal dendrites^32^, and dendritic CAMKIIα synthesis is required for the long-term maintenance of LTP and memory^33^. To examine the effect of FTLD-tau mutants on activity-dependent synthesis of CAMKIIα, we used a reporter containing the 5’ and 3’ untranslated regions (UTRs) of CAMKIIα fused with the coding sequence of myristoylated destabilized GFP (myr-dGFP-CamKIIα UTR)^32^. The myristoylation tag restricts diffusion of the GFP reporter to enable monitoring of local protein synthesis in neurons^32,47^. Myr-dGFP-CamKIIα UTR was co-transfected with mApple to delineate dendrite morphology in cultured rat hippocampal neurons that were either unstimulated or treated with cLTP, and myr-dGFP levels were measured in dendrites 4 hours post-LTP induction. After cLTP in control neurons, myr-dGFP expression was significantly increased in dendrites of neurons compared to unstimulated neurons (Fig. 2g,h). Neurons that were co-transfected with either tauV337M or tauP301S constructs did not exhibit an activity-dependent increase in reporter expression (Fig. 2g,h), indicating that both FTLD-tau mutants suppressed translation after LTP induction. Together, these data reveal that pathogenic FTLD-tau inhibits activity-induced protein synthesis in dendrites which is critical for LTP expression at synapses.

### FTLD-tau alters the LTP-associated translatome

To identify translated mRNAs that are dysregulated by tau during plasticity, we investigated the effect of FTLD-tau on the translatome in human neurons after LTP induction. Neuronal activity and plasticity induction leads to an increase in polysome-bound mRNAs associated with enhanced translation^44,48–50^. Thus, we used polysome profiling to isolate polysome-bound mRNAs for LTP-associated translatome analyses. TauWT and tauV337M_iso_ human neurons first received cLTP treatment, and polysome profiling was performed 45 min later followed by the extraction of RNA from the polysome fractions (Fig. 3a). The polysome peaks were significantly reduced in tauV337M_iso_ compared to tauWT neurons after cLTP induction (Fig. 3b,c), which is consistent with inhibited activity-dependent protein synthesis in tauV337M_iso_ neurons (Fig. 2c,d). RNA-seq of the polysome-bound mRNAs was performed to compare the abundance of translated mRNAs in tauV337M_iso_ and tauWT neurons (Extended Data Fig. 3a). Gene ontology (GO) enrichment analyses of the most downregulated transcripts in the translatome of tauV337M_iso_ neurons (1093 transcripts with log_2_FC < −0.5) uncovered major classifications as networks in dendrite and synaptic membrane compartments, as well as in the nucleus and nuclear lumen (Fig. 3d and Supplementary Table 1). The most upregulated polysome-bound transcripts in tauV337M_iso_ neurons (1225 transcripts with log_2_FC > 0.5) included β2M and other MHC-I complex proteins (Extended Data Fig. 3b and Supplementary Table 1), which is consistent with the upregulation of MHC-I expression during AD-related pathogenesis^51^.

**Fig. 3.**
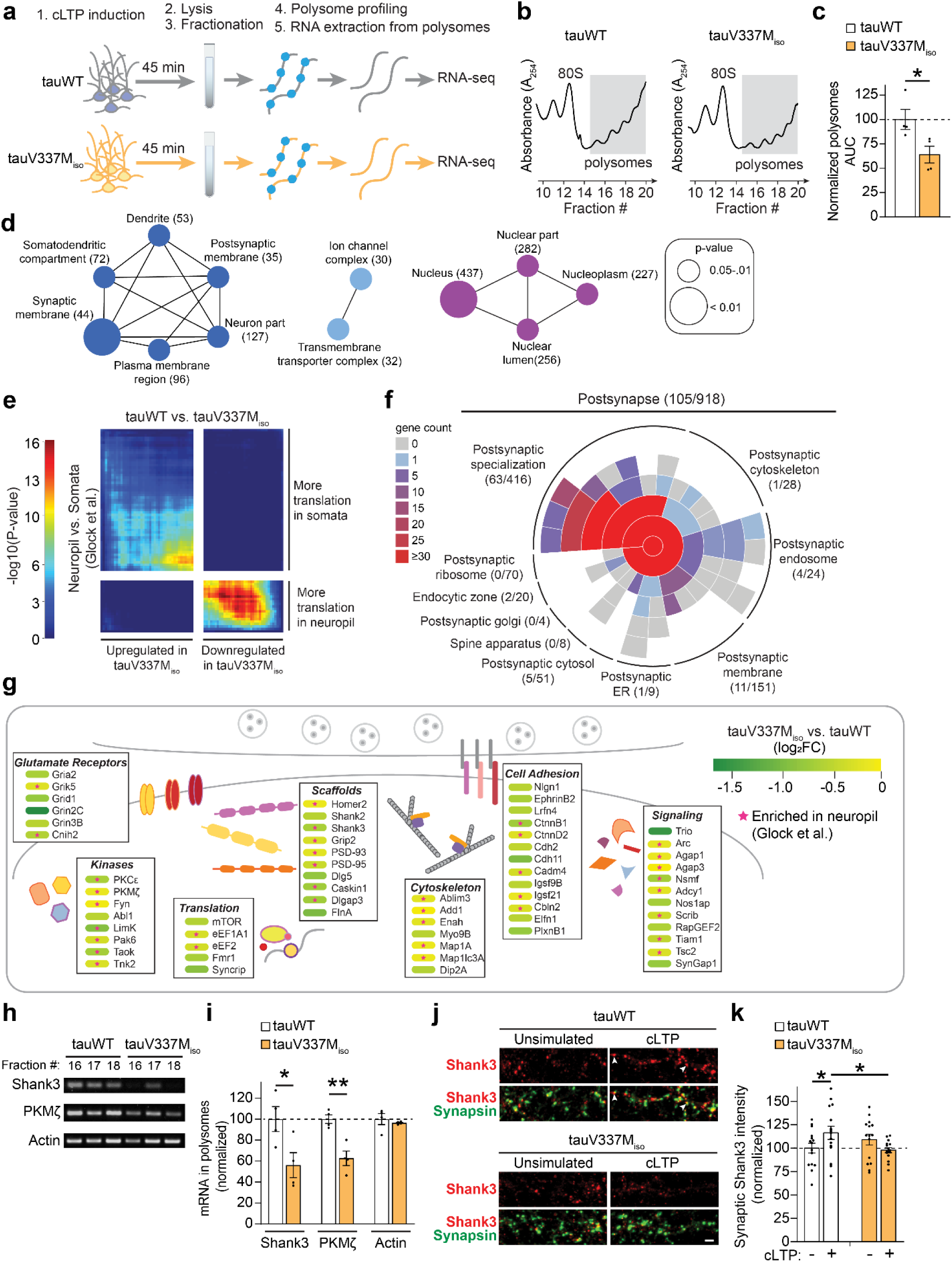
FTLD-tau mediated downregulation of synaptic mRNAs in the LTP-associated translatome. (**a**) Workflow of protocol to collect mRNAs extracted from polysomes in human neurons after cLTP induction. RNA-seq was performed to quantify the abundance of each transcript in polysomes extracted from tauWT and tauV337M_iso_ neurons (n = 4 cultures/group). (**b**) Representative polysome profiles from tauWT and tauV337M_iso_ neurons with the polysome peaks shaded in grey. (**c**) Quantification of the area under the curve (AUC) of the polysome peaks normalized to tauWT neurons (n = 4 cultures/group; *, p < 0.05, Student’s *t*-test). (**d**) ClueGO cellular component pathway enrichment of the 1,093 transcripts extracted from polysomes that were downregulated in tauV337M_iso_ neurons with < −0.5 log_2_FC compared to tauWT neurons. The number of downregulated transcripts detected within each GO term is indicated in parentheses. Node colors denote functionally grouped networks. (**e**) RRHO plot of the correlation between mRNAs translated in neuropil compared to somata^52^ and the differential LTP-associated translatome in tauV337M_iso_ compared to tauWT neurons. Analyses show a significant correlation in transcripts that were translated more within neuropil and downregulated in the tauV337M_iso_ LTP-associated translatome. (**f**) SynGO analyses of polysome-bound mRNAs that were downregulated in tauV337M_iso_ neurons and classified in the Postsynapse term. Analyses were performed on (**d**) transcripts with < −0.5 log_2_FC and (**e**) downregulated polysome-bound transcripts that were significantly correlated with more translation in neuropil^52^. (**g**) Select downregulated polysome-bound transcripts in tauV337M_iso_ neuron that were classified in the Postsynapse SynGO term (**f**). Color indicates log_2_FC of transcript abundance in polysomes of tauV337M_iso_ compared to tauWT neurons. Transcripts that are translated more within neuropil than in the soma of neurons^52^ are labeled by a star. (**h**) Representative RT-PCR results from three different fractions containing polysomes (#16-18) that were extracted from one culture of tauWT or tauV337M_iso_ neurons. (**i**) RT-PCR analyses of Shank3, PKMζ, and Actin mRNA levels in the polysome fractions (#16-18) that were each normalized to the mRNA levels in tauWT neurons (n = 4 cultures/group; * p < 0.05, **p<0.01, Student’s t-test). (**j**) Confocal images of Shank3 (red) and Synapsin (green) immunolabeling of tauWT and tauV337M_iso_ neurons with or without cLTP induction. Shank3 was increased at synapses after cLTP in tauWT neurons (arrows). Scale bar, 2 µm. (**k**) Quantification of synaptic Shank3 immunolabeling intensity co-localized with Synapsin (n = 15 images/group; * p < 0.05, two-way ANOVA, Bonferroni post-hoc analyses). Values were normalized to the mean Shank3 intensity at synapses in unstimulated tauWT neurons. Values are given as means ± SEM.

While many mRNAs have been detected within dendrites, the microdissection of neurons from rat hippocampus revealed distinct subcellular translatomes in the somatic compartment compared to the neuropil, consisting of dendrites and axons^52^. Using Rank-rank Hypergeometric Overlap (RRHO) analyses, we tested the relationship between the altered LTP-associated translatome in FTLD-tau neurons and the subcellular region-specific translatomes that were identified in neurons^52^. Interestingly, there was a striking correlation of 488 transcripts with both reduced translation in tauV337M_iso_ neurons and more translation in neuropil (Fig. 3e and Supplementary Table 1), highlighting the collection of downregulated translated mRNAs in tauopathy neurons within the neuropil-enriched translatome. Synaptic GO (SynGO) analyses of the transcripts with the lowest translation levels in tauV337M_iso_ neurons, including those that correlated with neuropil-enriched translation, classified 105 of the transcripts as postsynapse and subclassified 60% of these transcripts as postsynaptic specialization (Fig. 3f), which contains postsynaptic density protein ontologies (Supplementary Table 1). Notably, the mRNAs encoded proteins with a range of functions within the postsynaptic density ontology including glutamate receptors, kinases, scaffolding proteins, cytoskeleton-associated proteins, and cell adhesion molecules that span the synaptic cleft (Fig. 3g and Supplementary Table 1). Consistent with AMPAR dysregulation, tauV337M_iso_ neurons exhibited less LTP-associated polysome-bound mRNAs for GluA2 (Gria2), cornichon-2 (Cnih2), an auxiliary subunit of AMPARs that regulates channel kinetics and surface expression of GluA1-containing AMPARs^53^, and Trio, a Rho guanine exchange factor (RhoGEF) that modulates postsynaptic AMPARs and LTP expression^54^. Kinases with less polysome-bound mRNAs in FTLD-tau neurons and roles in modulating synapse strength included PKMζ^55^, PKCε^56^, LimK-1^57^, Abl1^58^, and Fyn^59^. Reduced mRNAs for scaffolding proteins included regulators of postsynaptic AMPARs and synapse structure such as PSD-93 and PSD-95^60^, DLG5^61^, GRIP2^62^, Caskin1^63^, Shank2^64^, and Shank3^65^. Shank3 is required for LTP^65^ and it regulates the exocytosis of GluA1-containing AMPARs at synapses^66^. RT-PCR was used to confirm that polysomes fractions from tauV337M_iso_ neurons after cLTP contained significantly lower levels of PKMζ and Shank3 mRNA compared to tauWT neurons (Fig. 3h,i). To monitor Shank3 protein levels at synapses of FTLD-tau neurons, tauWT and tauV337M_iso_ neurons were fixed 24 hours after cLTP induction for Shank3 immunolabeling. Postsynaptic Shank3 levels were significantly increased in tauWT neurons after cLTP compared to unstimulated neurons, but the cLTP-induced increase of postsynaptic Shank3 was blocked in tauV337M_iso_ neurons (Fig. 3j,k), confirming that activity-dependent changes in synaptic Shank3 were inhibited by FTLD-tau. Together, these results support that, following LTP induction, FTLD-tau inhibits the translation of a subset of mRNAs encoding postsynaptic regulators of synapse strength and plasticity.

### Increased FTLD-tau binding to eIF4B coincides with deficient eIF4B in the translation initiation complex

Mapping of the tau interactome in human neurons revealed eIF4B as a new tau-binding protein^11^, which led us to further examine eIF4B because of its function in activity-dependent translation in neurons^67,68^. To validate and probe tau-eIF4B binding in cells, we used a proximity ligation assay (PLA)^69^. PLA provides the unique advantage that it labels protein-protein interactions in individual cells retaining their intrinsic structures and cellular components, which is distinct from the co-immunoprecipitation method that assays protein complexes detected in homogenized lysates of cells. We first used PLA with HA and flag antibodies on HEK293 cells co-expressing GFP with HA-tagged eIF4B and either flag-tagged tauWT or tauV337M, and we quantified the mean PLA intensity to evaluate the protein-protein interactions in each cell. Cells co-expressing eIF4B and either tauWT or tauV337M had significantly greater PLA signal than control cells that expressed only eIF4B or tauV337M (Fig. 4a,b), confirming that both tauWT-flag and tauV337M-flag interacted with HA-eIF4B in HEK293 cells.

**Fig. 4.**
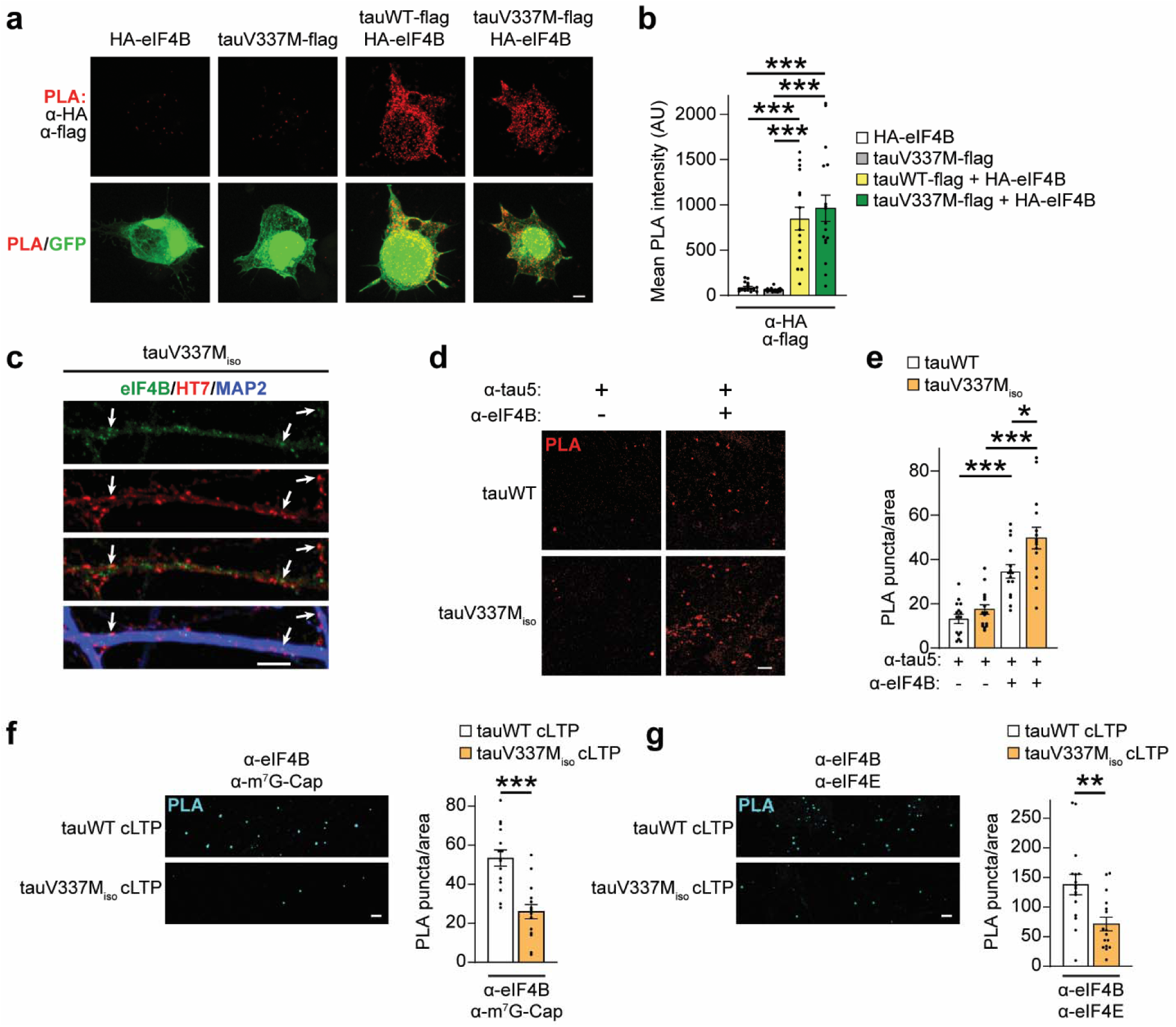
Enhanced binding of tau to eIF4B in tauV337M neurons coincides with dissociation of eIF4B from translation initiation complex components. (**a**) Images of HEK293 cells that were co-transfected with GFP (green), HA-tagged human eIF4B, and flag-tagged human tau with or without the V337M mutation. Labeling by the proximity ligation assay (PLA) with anti-HA and anti-flag antibodies marks the binding of eIF4B and tau (red). Controls for the PLA specificity using these antibodies included HEK293 cells expressing HA-eIF4B or tauV337M-flag alone. Scale bar, 5 µm. (**b**) Graph of the mean PLA signal intensity measured in HEK293 cells co-expressing HA-eIF4B with tauWT-flag or tauV337M-flag, and control HEK293 cells expressing HA-eIF4B alone and tauV337M-flag alone (AU: arbitrary units; n = 15-22 cells/group; *** p < 0.001, two-way ANOVA, Bonferroni post-hoc analyses). (**c**) Images of dendrites labeled by MAP2 staining (blue) in tauV337M_iso_ neurons in which co-localization of tau and eIF4B (arrows) was evident by immunostaining with HT7 (red) and eIF4B (green) antibodies, respectively. Scale bar, 5 µm. (**d**) PLA of the interaction between endogenous eIF4B and tau (red) using anti-eIF4B and anti-tau5 antibodies on tauWT and tauV337M_iso_ neurons. Scale bar, 5 µm. (**e**) Quantification of the eIF4B-tau PLA puncta density in tauWT and tauV337M_iso_ human neurons compared to negative controls with only anti-tau5 labeling (n = 15 images/group; * p < 0.05, *** p < 0.001, one-way ANOVA, Bonferroni post-hoc analyses). PLA puncta density was quantified for each 101 µm × 101 µm image. (**f** and **g**) Representative images and quantification of the PLA signal in human neuron cultures after cLTP induction detecting the interaction between (**f**) eIF4B and the 5’ m^7^G-Cap on mRNAs in the translation initiation complex (n = 15 images/group; *** p < 0.001, Student’s *t*-test) and between (**g**) eIF4B and eIF4E, the 5’ mRNA m^7^G-Cap binding protein (n = 16 images/group; ** p < 0.01, Student’s *t*-test). PLA puncta density was quantified for each 232 µm × 232 µm image. Scale bar, 5 µm. Values are given as means ± SEM.

In tauV337M_iso_ human neurons, eIF4B was detected in dendrites where it co-localized with HT7 immunolabeling of tau protein (Fig. 4c). To directly probe the interaction of eIF4B with tau in human neurons, the PLA signal from staining with eIF4B and tau5 antibodies was measured. TauV337M_iso_ neurons exhibited significantly greater PLA signal than tauWT neurons (Fig. 4d,e). Increased PLA signal was also detected in tauV337M_pat_ neurons compared to tauV337V_iso_ neurons (Extended Data Fig. 4a,b), confirming that eIF4B interacts more with FTLD-tau than with wild-type tau protein in neurons. We next examined whether the associations of eIF4B with the translation initiation complex and capped mRNA were altered by FTLD-tau. The eIF4F complex, comprised of the mRNA cap-binding protein eIF4E, the scaffold protein eIF4G, and the helicase eIF4A binds to mRNA with a 5’ m^7^G-Cap for translation initiation. We performed PLA on human neurons after cLTP induction with α-eIF4B and either α-m^7^G-Cap or α-eIF4E to assess the association of eIF4B with the translation initiation complex during LTP expression. The density of α-eIF4B and α-m^7^G-Cap PLA signal was significantly reduced in tauV337M_iso_ compared to tauWT neuron cultures (Fig. 4f). Likewise, α-eIF4B and α-eIF4E PLA signal was significantly reduced in tauV337M_iso_ neurons (Fig. 4g), suggesting that FTLD-tau inhibits the interaction between eIF4B and the translation initiation complex. To confirm whether the eIF4F components in the initiation complex remain intact in tauV337M_iso_ neurons, we performed PLA using α-eIF4E or α-eIF4G with α-m^7^G-Cap antibodies, which showed that FTLD-tau did not affect binding of eIF4E or eIF4G to the 5’ m^7^G-Cap of mRNAs (Extended Data Fig. 4c,d). These findings suggest that binding of pathogenic tau to eIF4B in neurons inhibits the function of eIF4B in the translation initiation complex.

### Deficient eIF4B in tauopathy models and human AD brain

To further examine the role of eIF4B in local translation dysregulation in tauopathy, we performed immunolabeling and quantified eIF4B levels in dendrites, marked by MAP2, and at synapses, marked by vGluT1, in FTLD-tau human neurons. Both dendritic and synaptic eIF4B levels were significantly decreased in tauV337M_iso_ compared to tauWT neurons (Fig. 5a-d). Nonetheless, levels of eIF4A, eIF4E and eIF4G were not significantly altered at synapses in tauV337M_iso_ neurons (Extended Data Fig. 5a-c), suggesting that the loss of eIF4B was not due to an overall deficiency in translation initiation complex proteins. A proteomics study of human brain tissues identified eIF4B in a co-expression module of proteins that were downregulated in AD brain and significantly correlated with pathology and cognition^70^. We assessed eIF4B by immunoblotting of post-mortem human brain tissue homogenates from the middle temporal gyrus of cases that were designated as either control (n = 7) or AD (n = 7, score A3B3C3) based on neuropathological evaluation (Fig. 5e and Supplementary Table 2). Total levels of soluble eIF4B were significantly reduced in AD brains compared to controls with no significant change in the levels of eIF4E (Fig. 5f). This links eIF4B deficiency also with AD-related pathogenesis, which has shared pathological features with FTLD-tau caused by *MAPT* mutations^71^.

**Fig. 5.**
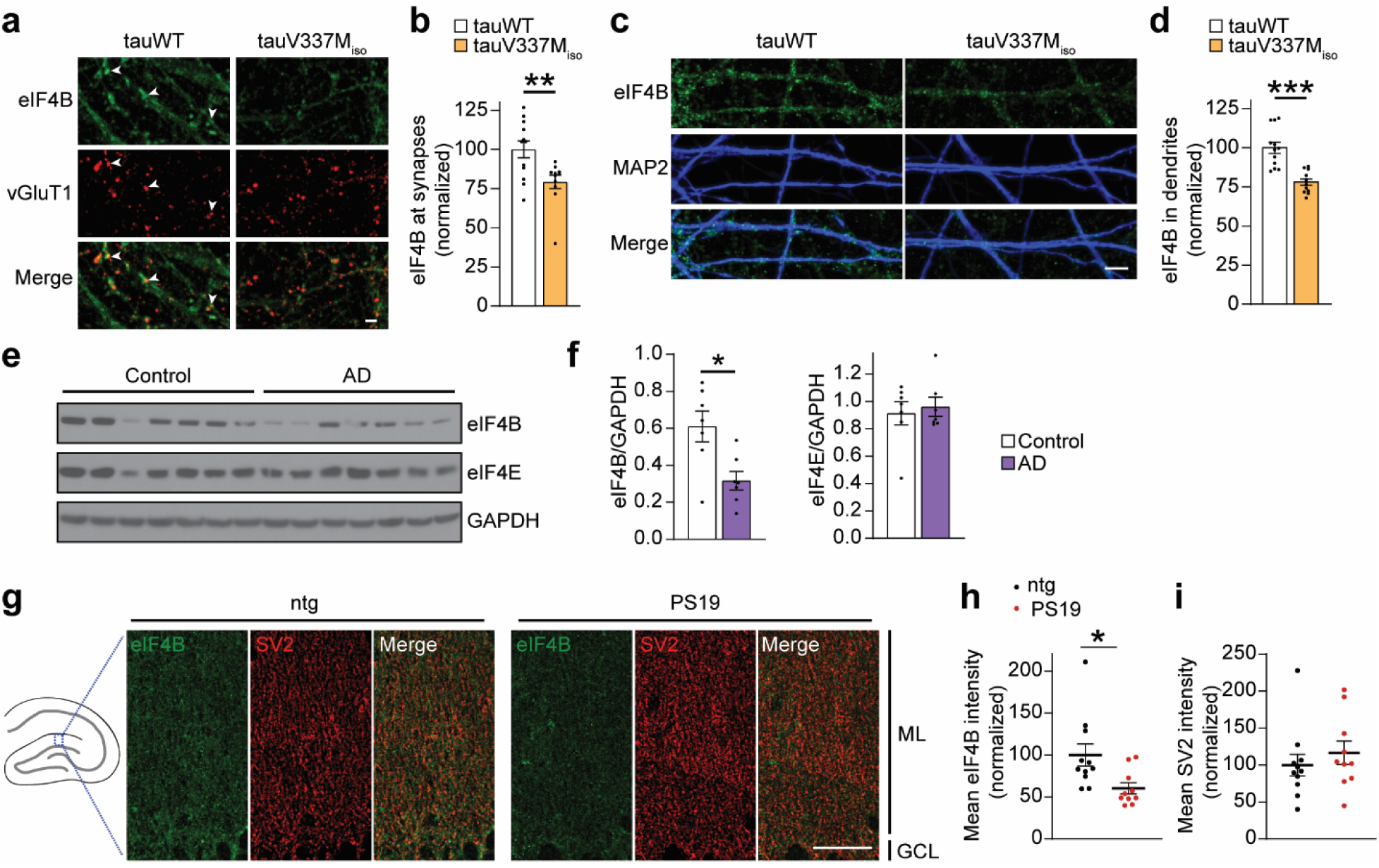
Reduced eIF4B levels in FTLD-tau human neurons, AD brain tissue, and the hippocampus of PS19 mice. (**a**) Images of immunolabeling for eIF4B (green) co-localized with vGluT1 puncta (red) at synapses in tauWT and tauV337M_iso_ human neurons. TauWT neurons had more eIF4B at synapses (arrowheads). Scale bar, 2 µm. (**b**) The mean intensity of eIF4B immunoreactivity in synapses marked by vGluT1 was quantified and normalized to the mean intensity of eIF4B in tauWT neurons (n = 11-12 images/group; ** p < 0.01, Student’s *t*-test). (**c**) Images of immunolabeling for eIF4B (green) co-localized with MAP2 (blue) in dendrites of tauWT and tauV337M_iso_ human neurons. Scale bar, 5 µm. (**d**) Quantification of eIF4B immunolabeling intensity in dendrites of tauWT and tauV337M_iso_ human neurons that were labeled by MAP2 immunofluorescence. Values were normalized to the mean intensity of eIF4B in tauWT neurons (n = 12 images/group; *** p < 0.001, Student’s *t*-test). (**e**) Immunoblots of RIPA-soluble homogenates from the middle temporal gyrus of human brains to detect total soluble protein levels of eIF4B and GAPDH. Control and AD cases were classified by neuropathological analyses. (**f**) Quantification of western blot immunoreactivity for soluble eIF4B and eIF4E levels in human brain tissue normalized to GAPDH levels (n = 7 samples/group; *, p < 0.05, Student’s *t*-test). (**g**) An illustration of the hippocampus depicts the region in the dentate gyrus that was imaged and analyzed (blue box). Confocal high magnification images of co-immunolabeling of eIF4B (green) and SV2 (red), a synaptic marker, were focused on synapses in the molecular layer (ML) of the dentate gyrus of 6-7-month-old non-transgenic (ntg) and PS19 mice with only the top of the dentate granule cell layer (GCL) visible. Scale bar, 20 µm. (**h, i**) Quantification of the mean intensities of (**h**) eIF4B and (**i**) SV2 immunofluorescence in the molecular layer of the dentate gyrus of 6-7-month-old non-transgenic (ntg) and PS19 mice (n = 10-11 mice/group; * p < 0.05, Student’s *t*-test). Values are given as means ± SEM.

We next investigated eIF4B in the PS19 mouse model of tauopathy, which has transgenic expression of human tau carrying the P301S mutation in neurons^2^. Hippocampal synapse pathophysiology begins as early as 3 months of age in PS19 mice, followed by neurodegeneration and the formation of pathological tau aggregates starting at 9 months old^2^. We examined immunolabeling of eIF4B in the dentate gyrus where protein synthesis is required for the sustained potentiation of postsynaptic strength during plasticity associated with memory consolidation in dentate granule cells (DGCs)^72^. The eIF4B levels in DGC dendrites of the molecular layer were significantly decreased in 6-7-month-old PS19 mice compared to littermate non-transgenic (ntg) mice (Fig. 5g,h). Immunostaining of SV2, a presynaptic marker, was not significantly different in the molecular layer of ntg and PS19 mice (Fig. 5g,i), suggesting that synaptic inputs remained intact in the dentate gyrus of 6-7-month-old PS19 mice.

In aged PS19 mice (> 9 months old), pathological tau co-aggregates with RNA-binding proteins in the brain such as TIA1^73,74^. Similarly, co-immunolabeling of eIF4B and phospho-threonine 231 tau (p-tau, AT180) in the hippocampus of 10-11-month-old PS19 mice revealed inclusions with overlapping p-tau and eIF4B reactivity within CA1 (Extended Data Fig. 5d). These results suggest that, after the downregulation of eIF4B in 6-7-month-old PS19 mice (Fig. 5g,h), eIF4B can also form co-aggregates with tau in older PS19 mice when they reach an advanced stage of neuropathology. Overall, our findings suggest that pathogenic tau drives the loss of eIF4B function in neurons, involving both the binding of tau to eIF4B and the downregulation of eIF4B levels in dendrites.

### eIF4B protects against FTLD-tau mediated LTP and memory impairments

We next tested the impact of eIF4B overexpression on pathophysiology in both human neuron and PS19 mouse tauopathy models. An empty vector control lentivirus (lenti-control) and a lentivirus for flag-tagged eIF4B expression (lenti-eIF4B) were generated. Expression of flag-tagged eIF4B and its localization in MAP2-labeled dendrites was confirmed in the infected human neurons (Fig. 6a). During differentiation, tauV337M_iso_ cells were infected with either lenti-control or lenti-eIF4B two days before neuron plating, and synaptic plasticity was evaluated in the neurons 6 weeks later. Consistent with our previous results demonstrating that FTLD-tau blocks LTP expression in human neurons, the surface staining of GluA1-containing AMPARs at synapses was not changed in cLTP-treated compared to unstimulated tauV337M_iso_ lenti-control neurons (Fig. 6b,c). TauV337M_iso_ lenti-eIF4B neurons exhibited significantly increased surface GluA1 immunolabeling at synapses following cLTP compared to both unstimulated controls and to tauV337M_iso_ lenti-control neurons with cLTP (Fig. 6b,c), indicating that increased eIF4B expression restored AMPAR trafficking during synaptic plasticity in FTLD-tau neurons. We next evaluated the effect of eIF4B expression on activity-dependent protein synthesis in dendrites of FTLD-tau neurons. As expected, the intensity of puromycin labeling in dendrites did not change after cLTP induction in tauV337M_iso_ lenti-control neurons compared to unstimulated controls, but in tauV337M_iso_ lenti-eIF4B neurons, cLTP induced a significantly greater intensity of puromycin labeling in dendrites compared to controls (Fig. 6d,e). Thus, increasing eIF4B levels in FTLD-tau human neurons was sufficient to reestablish the activity-induced local protein synthesis that is required for the trafficking of AMPARs during LTP expression.

**Fig. 6.**
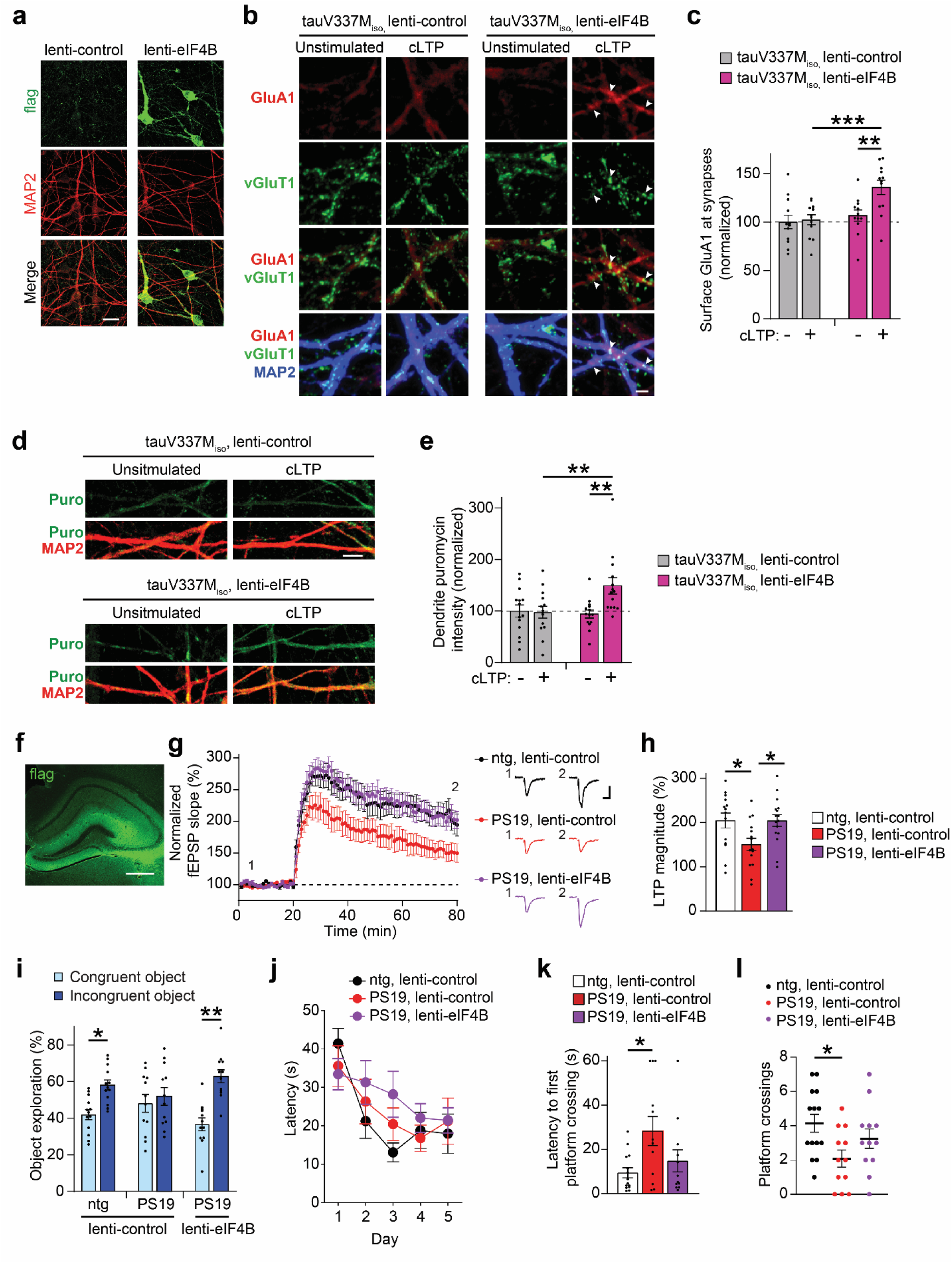
Enhancing eIF4B levels restores plasticity and memory in human and mouse tauopathy models. (**a**) TauV337M_iso_ neurons were infected with a control empty vector lentivirus (lenti-control) or a flag-tagged eIF4B lentivirus (lenti-eIF4B) and immunostained for anti-flag (green) to label exogenously expressed eIF4B and anti-MAP2 (red) to label dendrites. Scale bar, 20 µm. (**b**) Images of surface GluA1 (red) co-localized with vGluT1 (green) at synapses on dendrites (blue) revealed increased postsynaptic AMPARs in lenti-eIF4B infected tauV337M_iso_ neurons after cLTP induction (arrowheads). Scale bar, 2 µm. (**c**) Surface GluA1 immunolabeling at synapses was quantified in tauV337M_iso_ neurons infected with lenti-control or lenti-eIF4B that were unstimulated or received cLTP treatment and values were normalized to the unstimulated tauV337M_iso_ lenti-control group (n = 10 images/group; ** p < 0.01, *** p < 0.001, two-way ANOVA, Bonferroni post-hoc analyses). (**d**) Images of anti-puromycin immunolabeling (green) of newly synthesized proteins in dendrites (red) of lenti-control or lenti-eIF4B infected tauV337M_iso_ neurons. Scale bar, 5 µm. (**e**) Quantification of mean puromycin immunostaining intensity in dendrites of tauV337M_iso_ neurons with or without eIF4B overexpression (n= 14 images/group; ** p < 0.01, two-way ANOVA, Bonferroni post-hoc analyses). All values were normalized to the mean puromycin intensity in dendrites of unstimulated tauV337M_iso_ lenti-control neurons. (**f**) Representative image of lentivirus-based expression of flag-tagged eIF4B (green) in mouse hippocampus. Scale bar, 500 µm. (**g-l**) Bilateral hippocampi of mice were injected with lenti-control or lenti-eIF4B at 5-6 months of age. For electrophysiology, acute slices were prepared from mice 6-8 weeks later for field recordings from the dentate gyrus. For behavior analyses, mice were injected with lentivirus at 6-7 months of age and subjected to tests 4-6 weeks later. (**g**) Field recordings were performed in dentate gyrus of acute brain slices with LTP induced by TBS after a 20 min baseline period. Scale bars: 0.5 mV and 10 ms. (**h**) Graph of the mean LTP magnitude at 55-60 min following TBS induction (n=14-16 from 5 mice/group, * p < 0.05, one-way ANOVA, Bonferroni post-hoc analyses). (**i**) The percentage of time that mice spent exploring the congruent and incongruent objects in the object-context discrimination test of pattern separation memory was analyzed (n= 12-14 mice/group, * p < 0.05, ** p < 0.01, paired Student’s *t*-test). (**j**) The latency to find the hidden platform in the Morris water maze test was not significantly different between groups (n = 12-14 mice/group, analyzed by repeated measures two-way ANOVA). (**k**) Mean latency to the first platform crossing and (**l**) mean total platform crossings in the 24 h probe test of spatial memory (n= 12-14 mice/group, * p < 0.05, one-way ANOVA, Bonferroni post-hoc analyses).

Lenti-eIF4B injection into the hippocampus of PS19 mice was performed to evaluate the impact of augmenting eIF4B levels *in vivo* on tauopathy-related pathophysiology and cognitive impairment. At 5-6 months of age, stereotaxic bilateral injection of lenti-control or lenti-eIF4B was targeted to the DGCs where synaptic eIF4B levels were downregulated in PS19 mice (Fig. 5g,h). Expression of flag-tagged eIF4B was confirmed in the molecular layer containing DGC dendrites (Fig. 6f). To assess basal synaptic transmission in brain slices from ntg lenti-control, PS19 lenti-control and PS19 lenti-eIF4B mice 6-8 weeks after lentivirus stereotaxic injection, field potentials in the molecular layer were elicited by perforant pathway excitation at ascending stimulus intensities. Input/output analyses showed no overall significant differences between groups, but PS19 lenti-control slices showed a moderate increase in fiber volley amplitude and basal postsynaptic responsiveness compared to ntg lenti-controls at a subset of stimulus intensities tested (Extended Data Fig. 6a). Theta burst stimulation (TBS) of the perforant pathway induced persistent LTP in DGCs of ntg lenti-control mice and PS19 lenti-eIF4B mice with a comparable magnitude of enhancement in synaptic strength (Fig. 6g,h). Conversely, the LTP magnitude was significantly reduced in PS19 lenti-control mice compared to both ntg lenti-control and PS19 lenti-eIF4B mice (Fig. 6g,h), indicating that eIF4B overexpression was sufficient to protect DGCs in PS19 mice from impaired synaptic plasticity.

Behavioral tests of hippocampus-dependent memory were conducted on mice at 7-8 months of age to evaluate the effect of eIF4B on tau-mediated memory loss. In the object-context discrimination test, the ntg lenti-control and PS19 lenti-eIF4B mice both showed intact pattern separation memory whereas the PS19 lenti-control mice demonstrated impaired memory (Fig. 6i and Extended Data Fig. 6b). In the Morris water maze test, PS19 mice displayed normal spatial learning during hidden platform training (Fig. 6j). During the probe test of spatial memory, PS19 lenti-control mice took significantly longer than ntg lenti-control mice to cross the location of the platform and had less platform crossings (Fig. 6k,l). On the other hand, PS19 lenti-eIF4B mice exhibited latency to first platform crossing and number of platform crossings results that were similar to ntg lenti-control mice without a change in swim speed (Fig. 6k,l and Extended Data Fig. 6c), suggesting that eIF4B overexpression improved spatial memory in PS19 mice. Together, these data support that increasing eIF4B levels in tauopathy neurons restores the protein synthesis in dendrites required for LTP expression in conjunction with protection against tauopathy-related memory loss in PS19 mice.

### Obstructing the FTLD-tau interaction with eIF4B prevents synapse dysfunction

To further investigate the physiological impact of the tau-eIF4B interaction, we identified the tau-interacting domain of eIF4B. A series of truncated eIF4B constructs were made (Extended Data Fig. 7a), and their interactions with tauV337M co-expressed in HEK293 were probed by PLA (Extended Data Fig. 7b). Interestingly, eIF4B Δ367-423 and Δ424-611 constructs, with deletions in the arginine rich motif (ARM) and the C-terminal domain (CTD), respectively, both showed significantly reduced binding to tauV337M compared to full-length eIF4B (Extended Data Fig. 7c). However, the aspartic acid, arginine, tyrosine, and glycine rich (DRYG) domain of eIF4B that binds to eIF3^75^ and the RNA recognition motif (RRM) domain were not required for the eIF4B-tau interaction (Extended Data Fig. 7c). We next created a construct containing only residues 351-611 of eIF4B (eIF4B_351-611_) with the ARM and CTD (Extended Data Fig. 7a) and examined its associations with tau and the translation initiation complex in HEK293 cells with GFP co-transfection. Using anti-HA and anti-flag PLA, we found that cells with HA-eIF4B_351-611_ or full-length HA-eIF4B both demonstrated significantly increased PLA signal with flag-tauV337M compared to control cells without eIF4B expression (Fig. 7a,b), indicating that eIF4B_351-611_ was sufficient to bind to tau. We next examined whether eIF4B_351-611_ was sufficient to interact with endogenous eIF4E in cells. The PLA intensity with anti-HA and anti-eIF4E antibodies was significantly increased in HEK293 cells expressing full-length HA-eIF4B compared to control cells (Fig. 7c,d), confirming eIF4B’s association with endogenous eIF4E. However, the PLA intensity was significantly decreased to control levels when HA-eIF4B_351-611_ was expressed in HEK293 cells (Fig. 7c,d), suggesting that eIF4B_351-611_ has a minimal interaction with the translation initiation complex.

**Fig. 7.**
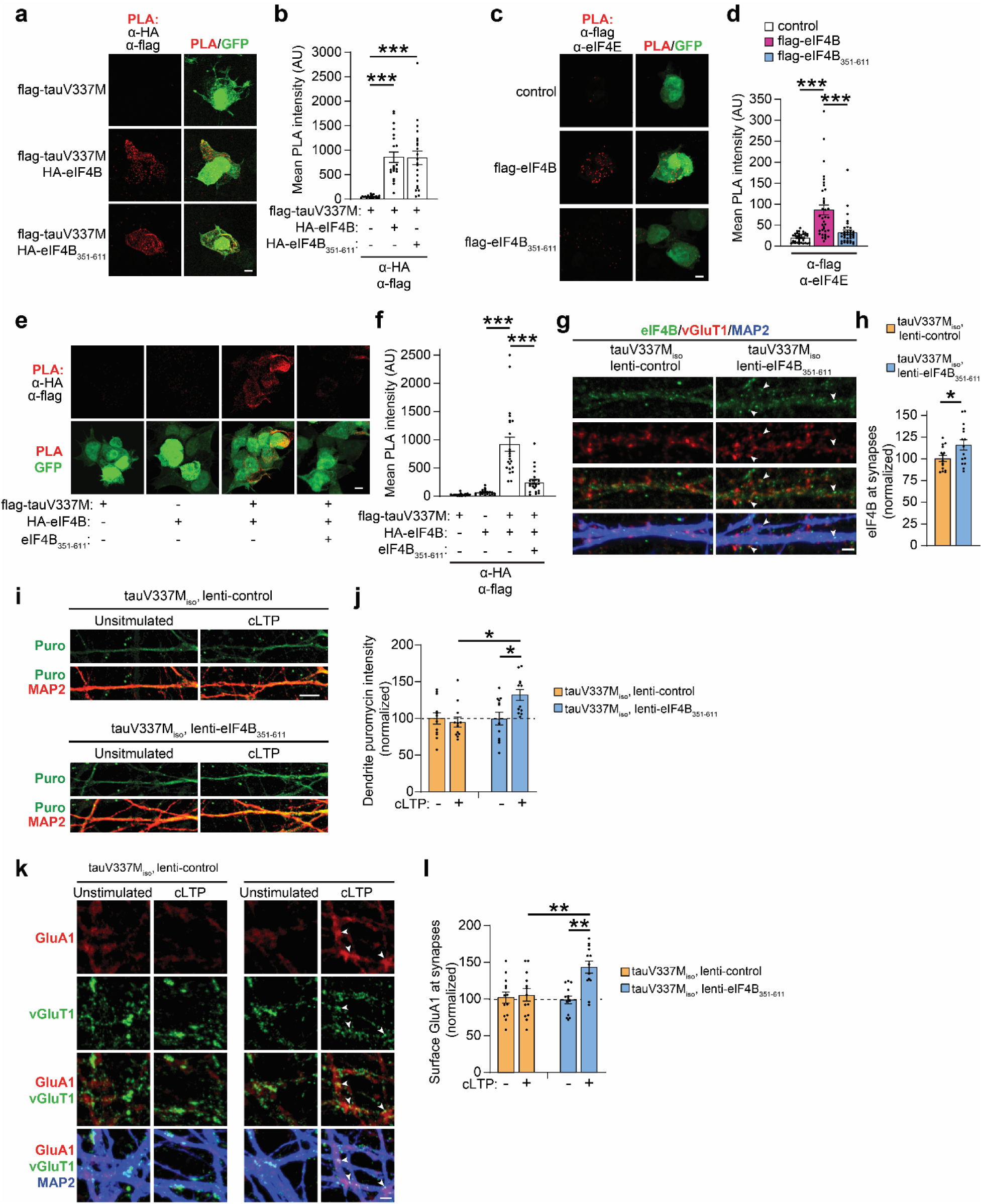
An inhibitor of the eIF4B and FTLD-tau interaction restores protein synthesis and postsynaptic AMPAR trafficking during LTP expression. (**a**) Confocal images of PLA signal (red) in HEK293 cells transfected with GFP (green) and flag-tagged tauV337M constructs alone or co-transfected together with either HA-tagged full-length eIF4B or HA-tagged eIF4B_351-611_. The anti-HA and anti-flag antibodies were used for PLA to label the flag-tauV337M and HA-eIF4B or HA-eIF4B_351-611_ in close proximity. Scale bar, 5 µm. (**b**) Quantification of the mean PLA intensity using anti-HA and anti-flag antibodies in HEK293 cells expressing flag-tauV337M with HA-tagged eIF4B constructs. (n = 21-22 cells/group; *** p < 0.001, one-way ANOVA, Bonferroni *post*-hoc analyses). (**c**) Confocal images of PLA signal (red) in HEK293 cells co-transfected with GFP (green) and flag-tagged full-length eIF4B or eIF4B_351-611_ and a control group with transfection of GFP alone. The anti-flag and anti-eIF4E antibodies were used for PLA to assess the interaction of the overexpressed flag-tagged full-length eIF4B or eIF4B_351-611_ fragment with endogenous eIF4E in HEK293 cells. Scale bar, 5 µm. (**d**) Quantification of the mean PLA intensity using anti-flag and anti-eIF4E antibodies in HEK293 cells expressing either flag-eIF4B or flag-eIF4B_351-611_ compared to control cells (n = 36 cells/group; *** p < 0.001, one-way ANOVA, Bonferroni *post*-hoc analyses). (**e**) Confocal images of PLA signal (red) in HEK293 cells co-transfected with GFP (green), flag-tauV337M, and HA-eIF4B with or without eIF4B_351-611_ expression. Control groups included HEK293 cells co-transfected with GFP and flag-tauV337M or HA-eIF4B alone. The anti-HA and anti-flag antibodies were used for PLA. Scale bar, 5 µm. (**f**) Quantification of the mean PLA intensity using anti-HA and anti-flag antibodies in HEK293 cells expressing flag-tauV337M and HA-full-length eIF4B with or without eIF4B_351-611_. Control groups included cells expressing either flag-tauV337M or HA-eIF4B alone (n = 21-23 cells/group; *** p < 0.001, one-way ANOVA, Bonferroni *post*-hoc analyses). (**g**) Confocal images of eIF4B (green) at synapses marked by vGluT1 (red) on dendrites (blue) in tauV337M_iso_ neurons infected with lenti-eIF4B_351-611_ or lenti-control. Endogenous eIF4B levels at synapses were increased in neurons with lenti-eIF4B_351-611_ (arrowheads). Scale bar, 2 µm. (**h**) Quantification of the mean eIF4B intensity at synapses in tauV337M_iso_ neurons infected with either lenti-control or lenti-eIF4B_351-611_. Analyses were normalized to the mean eIF4B intensity at synapses of tauV337M_iso_ lenti-control neurons (n = 14 images/group; * p < 0.05, Student’s *t*-test). (**i**) Confocal images of puromycin immunolabeling (green) of newly synthesized proteins in MAP2-labeled dendrites (red) of tauV337M_iso_ neurons that were infected with either lenti-eIF4B_351-611_ or lenti-control. Puromycin labeling was performed on unstimulated neurons and neurons that were treated for cLTP induction. Scale bar, 5 μm. (**j**) Quantification of puromycin immunolabeling intensity in dendrites of tauV337M_iso_ neurons infected with lenti-eIF4B_351-611_ or lenti-control (n = 12 images/group; * p < 0.05, two-way ANOVA, Bonferroni post-hoc analyses). All values were normalized to the mean puromycin intensity in dendrites of the unstimulated tauV337M_iso_ lenti-control neurons. (**k**) Images of surface GluA1 (red) co-localized with vGluT1 (green) at synapses on dendrites (blue) of tauV337M_iso_ neurons infected with lenti-eIF4B_351-611_ or lenti-control. Surface GluA1 intensity was increased at synapses in tauV337M_iso_ lenti-eIF4B_351-611_ neurons after cLTP induction (arrowheads). Scale bar, 2 µm. (**l**) Analyses of surface GluA1-containing AMPARs in unstimulated or cLTP-treated tauV337M_iso_ neurons that were infected with lenti-eIF4B_351-611_ or lenti-control (n = 14-15 images/group; ** p < 0.01, two-way ANOVA, Bonferroni post-hoc analyses). Values were normalized to the surface GluA1 intensity at synapses on unstimulated tauV337M_iso_ lenti-control neurons.

Given that eIF4B_351-611_ binds to tau without reconstituting the eIF4B-eIF4E association for translation initiation, we tested whether eIF4B_351-611_ could serve as a direct inhibitor of the tau-eIF4B interaction. The interaction between full-length eIF4B and tauV337M was probed using PLA with anti-HA and anti-flag antibodies in HEK293 cells co-expressing HA-eIF4B and flag-tauV337M with or without untagged eIF4B_351-611_. Strikingly, the high PLA intensity detected in cells with HA-eIF4B and flag-tauV337M was significantly reduced to control levels by co-expression of eIF4B_351-611_ (Fig. 7e,f), revealing that eIF4B_351-611_ is a potent inhibitor of the FTLD-tau-eIF4B interaction. We next examined whether inhibiting the FTLD-tau-eIF4B interaction alters synaptic eIF4B levels in tauV337M_iso_ neurons. Neurons were infected with an eIF4B_351-611_ lentivirus (lenti-eIF4B_351-611_) and eIF4B_351-611_ was detected in the soma and dendrites (Extended Data Fig. 7d). We used an antibody raised against the N-terminal domain of eIF4B to quantify only endogenously expressed eIF4B in tauV337M_iso_ neurons infected with either lenti-eIF4B_351-611_ or lenti-control. We found that tauV337M_iso_ lenti-eIF4B_351-611_ neurons had significantly more endogenous eIF4B at synapses compared to tauV337M_iso_ lenti-control neurons (Fig. 7g,h) showing that the synaptic localization of eIF4B in neurons can be protected by inhibiting the interaction between eIF4B and FTLD-tau.

To evaluate whether blocking the interaction of FTLD-tau with eIF4B influences plasticity, we analyzed the effect of eIF4B_351-611_ expression on activity-dependent translation and LTP in tauopathy neurons. Dendritic puromycin labeling of newly synthesized proteins was significantly increased after cLTP induction in tauV337M_iso_ lenti-eIF4B_351-611_ neurons compared to both unstimulated neurons and cLTP-treated tauV337M_iso_ lenti-control neurons (Fig. 7i,j). Furthermore, lenti-eIF4B_351-611_ expression increased postsynaptic GluA1 insertion following cLTP induction in tauV337M_iso_ neurons (Fig. 7k,l). Therefore, the interaction between pathogenic tau and eIF4B causes LTP impairment in human neurons, and preventing this protein-protein interaction restores local protein synthesis and LTP expression at synapses.

## Discussion

LTP impairment is a critical pathophysiological change in neuronal function that underlies memory loss in neurodegenerative diseases. Our findings show that pathogenic tau inhibits LTP expression by blocking translation in dendrites, effectively disabling the synthesis of proteins that enhance synapse strength during plasticity. Translation initiation is the first step in protein synthesis, and neuronal activity during LTP induction stimulates translation initiation^76^. Interestingly, we found that pathogenic tau acts as an inhibitor of the translation initiation machinery during LTP through its binding to eIF4B. This reveals a direct mechanism by which tau blocks the complex cascade of postsynaptic events that couple LTP induction with LTP expression.

Consistent with the reported role for eIF4B in activity-dependent translation in neurons^67,68^, we showed that LTP-induced protein synthesis in dendrites was inhibited in FTLD-tau neurons with deficient dendritic eIF4B. Further, we identified plasticity-associated mRNAs downregulated in the LTP-associated translatome of FTLD-tau neurons, including previously identified mRNAs locally translated in neuropil of mouse hippocampus^52^. This is consistent with a model in which pathogenic tau blocks synaptic plasticity by inhibiting the translation of the subset of mRNAs in dendrites regulated by eIF4B. PKMζ mRNA, which was downregulated in the FTLD-tau neuron LTP-associated translatome, is an example from this subset that is dendritically translated^52^ and regulated by eIF4B^67^. While eIF4B has been implicated in the translation of mRNAs with highly structured 5’ UTRs, the full subset of translated mRNAs regulated by eIF4B and activity in neurons is unclear. Notably, we did not detect changes in basal translation in the dendrites or soma of FTLD-tau neurons. However, using a long duration to label newly synthesized proteins (> 4h), reduced basal translation was detected only in neurons that harbored pathological tau aggregates in K3 and rTg4510 tauopathy mouse models^46^. Reduced translation was also detected in cultured cells overexpressing tau, which was linked to tau’s association with ribosomal proteins^45^. Tau may block activity-dependent translation and basal translation through distinct molecular mechanisms depending on the extent of pathological tau accumulation. Indeed, multiple mechanisms could affect translation throughout the progression of tauopathy. Before neurodegeneration, the enhanced translation of a subset of mRNAs in a *Drosophila* tauopathy model was linked to increased nonsense-mediated mRNA decay^77^. Transcriptomic changes reported across human tauopathy models may also influence protein synthesis and neuronal pathophysiology^78,79^. Importantly, the mechanistic regulation of activity-dependent translation in neurons is uniquely critical for maintaining precise control of synaptic strength during plasticity, and it can be distinct from other signaling mechanisms that regulate basal translation^80,81^. Thus, our findings point toward a specific effect of tau on activity-dependent translation in dendrites through eIF4B dysregulation that aligns with the emergence of LTP impairment at synapses.

We discovered that tau interacts with the C-terminal region of eIF4B, thereby preventing the integration of eIF4B into the translation initiation complex. The increased tau-eIF4B interaction in FTLD-tau neurons occurred in parallel with reduced levels of eIF4B at synapses, suggesting a dual impact of pathogenic tau that drives eIF4B loss of function. Surprisingly, the decrease in eIF4B levels was not associated with changes in the abundance of other important translation initiation factors, including eIF4A, eIF4E and eIF4G, pointing towards a direct effect of tau on eIF4B. Intriguingly, an in-depth proteomics analyses of human AD brain tissues revealed a co-expression module comprised of eIF4B and cytoskeletal proteins that was downregulated in AD brains compared to controls^70^, suggesting that the loss of eIF4B may involve cytoskeletal dysregulation. Previous work on neuron cultures showed that the phosphorylation of eIF4B also regulates its localization at synapses^67^. Whether eIF4B phosphorylation modulates the effects of pathogenic tau on local translation and eIF4B localization or function during LTP is unknown. Moreover, it remains to be determined how the many possible forms of pathogenic tau may contribute to eIF4B downregulation. Although comprehensively characterizing the landscape of pathogenic tau species that can emerge during pathogenesis is challenging, a previous study showed that human iPSC-derived neurons with the V337M *MAPT* mutation accumulate tau oligomers and tau with an abnormal conformation (MC1-positive) by 2 months post differentiation^82^. Accordingly, these aberrant forms of tau accumulate at around the same time as LTP expression is impaired and eIF4B is downregulated in the tauV337M neurons.

Interestingly, we found co-aggregates of eIF4B and phosphorylated tau in aged 10-11-month-old PS19 mouse hippocampus, raising the possibility that eIF4B may become incorporated into stress granules at an advanced stage of neuropathology. Previous studies showed that pathological tau associates with other RNA-binding proteins in aggregates, including TIA1 and G3BP^73,83^. These interactions can promote stress granule formation and exacerbate tau aggregation, which is related to the co-localization of stress granules and tau inclusions in more severe disease states^73,84^. Aggregates of eIF4B are also co-localized with amyloid-β plaques in a familial AD mouse model^85^. However, we found that the LTP impairment and eIF4B deficiency in PS19 mice occurs at a younger age before eIF4B- and p-tau-containing aggregates emerged in older mice, indicating that the effect of tau on eIF4B function precedes the formation of pathological aggregates.

Pathogenic tau accumulates in the brain and contributes to cognitive decline across neurodegenerative diseases classified as tauopathies. LTP impairment is also a hallmark pathophysiology that emerges across different models of tauopathies, including AD and FTLD mouse models^2–4,6^. It will be important to explore how the tau-mediated inhibition of activity-dependent protein synthesis may contribute to pathophysiology and memory loss across models of different types of tauopathy, particularly those in which memory-related brain regions are impacted. Considering that synapse repair can promote functional resilience to tau toxicity^86^, blocking the interaction between pathogenic tau and eIF4B could serve as a potential therapeutic avenue to restore synapse function and ameliorate tauopathy-related memory loss. As a targeted intervention, this strategy would directly modify the synaptic mechanisms at the source of tau-mediated dysfunction.

## Acknowledgements

We thank Dr. Gary Scott, Dr. Pejmun Haghighi, the Buck Institute Mouse Phenotyping Core, the Buck Institute Morphology and Imaging Core, Lee Cohen-Gould and the Weill Cornell Medicine Microscopy and Image Analysis Core, Ethan Ellis, Aana Hahn and the Mount Sinai Genomics Core, and the Mount Sinai Center for Bioinformatics for research support, and Mary Redwine for administrative support. This work was supported by the NIH (R01 AG070193 and K01 AG057862 to T.E.T, U54 NS100717 and R01 AG079291 to L.G., P01 AG066591 to E.V.), the Alzheimer’s Association (AARFD-19-616386 to G.K.), an NIH T32 training grant (T32 AG00026624 to K.A.P-N) and the Larry L. Hillblom Foundation (to L.Y.). Support to generate iPSCs was provided by the Knight Alzheimer Disease Research Center at Washington University (NIH P30 AG066444, P01 AG03991, P01 AG026276), NIH K01 AG046374 (CMK), and the Rainwater Charitable Organization. Human tissue samples were provided by the Neurodegenerative Disease Brain Bank at the University of California, San Francisco, which receives funding support from NIH grants P01 AG019724 and P50 AG023501, the Consortium for Frontotemporal Dementia Research, and the Tau Consortium.

## Author Contributions

G.K. and T.E.T. conceived the project and designed experiments. G.K., D.L., I. L.W., V.S., D.S., Y.Y.N, J.H.C., K.A.P-N., and T.E.T. performed experiments. G.K., D.L., K.S., D.S., K.A.P-N., and T.E.T. analyzed data. L.G., C.K., W.W.S., L.T.G., S.S. and A.L.N. provided experimental models and tissues. I.L.W., J.H.C, Y.Y.N, L.Y., K.S., E.V. and D.F. provided technical support. T.E.T and G.K. wrote the manuscript.

## Supplementary Information

Extended Data Fig. 1-7

Supplementary Table S1. Excel file containing data related to Fig. 3 and Extended Data Fig. 3.

Supplementary Table S2. Information on the human brain tissues used for this study.

## Methods

### Cell Cultures

The generation of tauWT iPSCs (WTC11) and CRISPR-edited isogenic tauV337M_iso_ iPSCs that were engineered with integration of the inducible Ngn2 transgene into the AAVS1 locus was previously described^37^. The generation of tauV337M_pat_ iPSCs (Line name: ND32951A.15) from a *MAPT* mutation carrier and the CRISPR-edited isogenic tauV337V_iso_ iPSCs (Line name: ND32951A.15ΔB06) with the *MAPT* mutation corrected was previously described^38^. To generate tauV337M_pat_ and tauV337V_iso_ iPSC lines with inducible Ngn2, we used AAVS1 TALEN pairs and a pUCM donor vector (kind gifts from Dr. Bruce Conklin, Gladstone Institutes) containing a Tet-On 3G tetracycline-inducible Ngn2 transgene cassette flanked with AAVS1 homology arms and puromycin resistance^36^. The iPSCs were dissociated, then transfected the AAVS1 TALEN pairs and the Ngn2 pUCM vector using a Human Stem Cell Nucleofector Kit 1 (Lonza) with the Nucleofector 2b Device (Lonza). The iPSCs were plated after transfection and puromycin (0.1-0.3 ug/mL) was added 24-48 hrs later to select for edited iPSCs. Media was regularly changed with puromycin included while colonies of individual clones grew. Colonies were picked to culture individual clonal iPSC lines. The iPSC clones used for experiments were confirmed for integration of the Ngn2 transgene cassette into one AAVS1 allele by PCR. Normal karyotyping was confirmed in the clonal iPSC lines used. All iPSC cultures were grown in E8 media in a cell culture incubator with 5% CO2 at 37°C, and ROCK inhibitor (Y-27632, 10 μM, Cayman chemicals) was used for iPSC cultures at low density. The use of human iPSCs in this study was approved by the Committee on Human Research at the Buck Institute.

To initiate the conversion of iPSCs to neurons, human iPSCs (1.5 ×10^6^) were plated onto each well of a 6-well plate coated with Matrigel (Corning) for 3 days of pre-differentiation in Knockout DMEM/F-12 media containing doxycycline (2 ug/mL, Sigma), N2 supplement (100x, Gibco), MEM non-essential amino acid solution (100x, Gibco), brain-derived neurotrophic factor (10 ng/mL, Peprotech), neurotrophin-3 (10 ng/mL, Peprotech) and ROCK inhibitor (Y-27632, 10 μM, Cayman chemicals). Cell media was exchanged daily with fresh media without ROCK inhibitor for the next two days. On the fourth day, pre-differentiated cells were dissociated with Accutase (STEMCELL Technologies, Inc.) and plated onto Matrigel-coated coverslips or tissue culture plates for the growth of neuron cultures in Neurobasal-A media (Gibco) containing B27 supplement (50x, Gibco), GlutaMAX (400x, Gibco), BDNF (10 ng/mL) and NT3 (10 ng/mL) with doxycycline (2ug/mL). Cultured rat astrocytes (ScienCell) were added to the neurons within 24 hours after plating the neurons to support their maturation. Half of the media was replaced on day 5-6 with supplemented Neurobasal-A media containing cytosine β-D-arabinofuranoside (Ara-C) without doxycycline. Half of the media was removed again on day 10-11, and MEM media containing glucose (27.7mM), NaHCO_3_ (2.4mM), B-27 supplement, L-glutamine (0.5 mM, Gibco), Ara-C and fetal bovine serum (5%) was added to the existing media up to four times the remaining volume. One third of the culture media was replaced with fresh MEM media once per week. Neurons were allowed to mature for at least 6-8 weeks before use for experiments. For polysome profile experiments, 8 × 10^6^ neurons were plated on a 10 cm tissue Matrigel-coated tissue culture plate co-plated with approximately 500,000 rat astrocytes. For immunocytochemistry experiments, 200,000 neurons were plated into each well of a 24-well plate with up to 30,000-40,000 cultured rat astrocytes added.

HEK293FT cells (Invitrogen) were maintained in DMEM media supplemented with 10% fetal bovine serum (Gemini), 200 mM L-Glutamine, 10 mM non-essential amino acids, and 100 mM sodium pyruvate. Cells were not passaged more than 10 times beyond the original stock. Lipofectamine 2000 (Invitrogen) was used for transfection of HEK293 cells with plasmids.

Hippocampi from day 22 embryonic rats were dissected from timed pregnant rats (Charles River). The neurons were dissociated from hippocampi after a 20 min incubation at 37° in 0.03% trypsin. The hippocampal neurons were plated onto poly-l-lysine (Sigma)-coated coverslips in a 24-well plate coated with Neurobasal media (Gibco) containing B27 supplement (Gibco) and Glutamax (400x, Gibco). The next day, half of the culture media was replaced with fresh media. Another half of the media was replaced with fresh media containing Ara-C at 7 days *in vitro* (DIV). At 11 DIV, half of the culture media was reserved, and the rest was used for Lipofectamine 2000 (Invitrogen) transfection of the hippocampal neurons with plasmids. The culture media containing transfection reagents was replaced the next day by the reserved media to maintain healthy neurons. Experiments were performed with the neurons at 15 DIV.

### Antibodies

The following primary antibodies were used: 7-methylguanosine (m7G)-Cap (RN016M, MBL), AT180 (MN1040, Thermo Fisher), eIF4A (sc-377315, Santa Cruz), eIF4B (3592T, Cell Signaling), eIF4E (9742S, Cell Signaling), eIF4E (sc-376062, Santa Cruz), eIF4G (2498S, Cell Signaling), FLAG (F1804, Sigma-Aldrich), GAPDH (MAB374, Sigma-Aldrich), GFP-conjugated 488 (A21311, Invitrogen), GluA1 (ABN241, Millipore), HA (H6908, Sigma-Aldrich), rabbit MAP2 (4542S, Cell Signaling), chicken MAP2 (nb 300 213, Novus Biologicals), puromycin (EQ0001, Kerafast), Synapsin (5297S, Cell Signaling), RPS6 (sc-74459, Santa Cruz), Shank3 (sc-377088, Santa Cruz), SV2 (University of Iowa DSHB), Tau5 (AHB0042, Thermo Fisher), HT7 (MN1000, Thermo Fisher), vGluT1 (MAB5502, Millipore), GluN1 (114 011, Synaptic Systems). The following secondary antibodies were used: anti-mouse Alexa Fluor 488 (A11029, Invitrogen), anti-mouse Alexa Fluor 546 (A11030, Invitrogen), anti-chicken Alexa Fluor Plus 647 (PIA32933, Invitrogen), anti-chicken Alexa Fluor Plus 546 (A-11040, Invitrogen), anti-rabbit Alexa Fluor 488 (A-11034, Invitrogen), anti-rabbit Alexa Fluor 647 (A-21245, Invitrogen), anti-mouse HRP (115-035-166, Jackson ImmunoResearch), anti-rabbit HRP (111-035-144, Jackson ImmunoResearch).

### Plasmids and Lentivirus

Flag-tagged or HA-tagged full-length human eIF4B (SinoBiological) was cloned into pcDNA3.1. Truncated eIF4B constructs were engineered with site-directed mutagenesis by Genscript, Inc. to generate HA-tagged Δ97-175, Δ214-327, Δ367-423, or Δ424-611 deletions in eIF4B. Deletions of eIF4B domains were confirmed by sequencing. Flag-tagged, HA-tagged and untagged eIF4B_351-611_ plasmids were subcloned in pcDNA3.1. Human wildtype tau (2N4R) tagged with flag in pcDNA3.1 was mutagenized by PCR and cloned to make tauV337M and tauP301S constructs. For expression in dissociated hippocampal neurons, pcDNA3.1 5’-myrdEGFP-3’ plasmid was a gift from Dr. Erin Schuman (Addgene plasmid #16075) and a pGW1-mApple plasmid (gift from Dr. Steve Finkbeiner) was used.

Plasmids for lentivirus packaging included Δ8.9 and VSV-G (Gladstone Institutes). The flag-tagged eIF4B and flag-tagged eIF4B_351-611_ sequences were each cloned into an FUGW2 lentiviral expression plasmid (Gladstone Institutes). To make lentivirus, HEK293 cells were split and plated into 15 cm culture plates. The next day a calcium phosphate transfection method was used to transfect the cells with Δ8.9, VSV-G, and the empty vector control FUGW2, or flag-eIF4B FUGW2, or flag-eIF4B_351-611_ FUGW2. The culture media was replaced with fresh media the following day, and lentivirus was harvested from the media twice at 24 hrs and 48 hrs later. The collected media was filtered (0.45 μm pore size), and sucrose gradient ultracentrifugation was used to purify the lentivirus. The resulting lentiviral pellet was resuspended in sterile PBS. A p24 Rapid Titer Kit (Takara Bio USA, Inc) was applied to estimate lentivirus titer.

### Chemical LTP

Chemical LTP (cLTP) experiments on human neurons were performed in extracellular solution (ECS) containing (in mM): 125 NaCl, 5 KCl, 25 HEPES, 1 NaH_2_PO_4_, and 11 Glucose, and 2.5 CaCl_2_ (all from Sigma Aldrich) with 0.5 μM tetrodotoxin (Abcam) and 1 μM strychnine (Sigma-Aldrich) at pH 7.4 and warmed to 37°C. For cLTP induction in cultured rat hippocampal neurons 100 μM picrotoxin (Caymen Chemicals) was added to ECS. For the experiment to test NMDAR-dependence, the ECS contained 100 μM APV (Abcam) to block NMDAR activity. Anisomycin (40 μM, Sigma-Aldrich) was added to the ECS for the experiment to test the dependence of LTP on protein synthesis. Neurons were washed once in ECS then incubated in either ECS alone as an unstimulated control group or with 300 μM glycine (Sigma-Aldrich) in ECS to induce cLTP. Human neurons were incubated for 15 min in glycine at 37°C to induce cLTP then labeled with anti-GluA1 antibody in ECS (MilliporeSigma) to monitor surface AMPARs for 30 min at 37°C. For the 4 h and 24 h cLTP experiments, ECS was replaced with original cell culture media and incubated at 37°C until fixation. Rat neurons were incubated with glycine for 5 min at 37°C for cLTP induction then washed with ECS for 25 min at 37°C. At the end of each incubation, neurons were washed twice with ECS and fixed in 4% PFA (Electron Microscopy Sciences) in PBS. Coverslips were washed three times in PBS then incubated in blocking solution containing PBS, 2% normal goat serum (Jackson ImmunoResearch), and 0.1% Triton X-100 for 1 h. Human neurons were labeled with mouse anti-vGluT1 (MAB5502, Millipore) antibody in blocking solution for 1 hour at room temperature. After three washes in PBS, coverslips were incubated in anti-mouse Alexa Fluor 488 (A11001, Invitrogen) and anti–rabbit Alexa Fluor 647 (A21245, Invitrogen) secondary antibodies mixed in blocking solution and applied for 1 hour at room temperature followed by three PBS washes. The integrated intensity of GluA1 signal was measured from at least 60 synapses per image using Fiji software (ImageJ). For puromycin labeling, neurons were incubated at 37°C for 30 min after cLTP induction then 10 μM puromycin was applied and neurons were incubated at 37°C for an additional 15 min. Coverslips were washed in ECS after puromycin labeling and then fixed in 4% PFA in PBS. The integrated intensity of puromycin staining was measured from at least 5 second-order dendrites as outlined by MAP2 co-staining per image. The sum of the puromycin integrated intensities was normalized to the sum of the total dendritic area outlined by MAP2 staining for each image. Fixed rat hippocampal neurons expressing myr-dGFP-CamKIIα UTR were incubated with anti-GFP 488 (A-21311, Invitrogen) to enhance the GFP reporter signal for quantification. Experiments examining the fluorescence intensity of myr-dGFP-CamKIIα UTR or eIF4B in dendrites were analyzed in a similar manner as puromycin. Images were acquired on a Zeiss LSM 700 or Nikon AX confocal microscope. Image acquisition settings were calibrated to keep the brightest pixels in dendrites slightly below saturation. Analyses were performed blind to the experimental conditions.

### Immunocytochemistry

Fixation of neuron cultures was performed in 4% PFA in PBS for 15 min then coverslips were washed with PBS three times for 5 min. Blocking solution containing PBS, 0.1% Triton X-100, and 2% normal goat serum was added to coverslips for 1 hour at room temperature. Neurons were labeled with primary antibodies mixed in blocking solution for 1 hour at room temperature then washed three times with PBS. Secondary antibodies were diluted in blocking solution and added to the coverslips for 1 hour at room temperature then washed in PBS. Coverslips were mounted on glass slides in Prolong Gold (Invitrogen). For GluN1 immunostaining, chilled methanol (−20°C) was used for fixation for 10 min at −20°C which was followed by washing with PBS three times for 5 min. Coverslips were incubated in blocking solution containing PBS, 0.1% Triton X-100, and 10% normal goat serum for 30 min, then incubated with primary antibodies for 2 hours in PBS with 3% normal goat serum at room temperature. The coverslips were washed three times in PBS for 5 min, incubated in secondary antibodies for 2 hours in PBS with 3% normal goat serum at room temperature, washed again with PBS three times and mounted on glass slides. Imaging was performed using a Zeiss LSM 700 or Nikon AX laser scanning confocal microscope with a 63x oil objective. Confocal z-sections were stacked by maximum intensity projections. Laser and gain settings for acquisition were set to minimize the saturation of pixel intensities. For representative images shown, the image contrast was uniformly adjusted identically for each channel within each experiment to facilitate the visualization of low-intensity signals. The intensity of fluorescence in images was quantified after background subtraction was applied to the entire image using Fiji software (ImageJ).

### Mice and Stereotaxic Surgery

To generate ntg and PS19 littermate mice for experiments, hemizygous human tauP301S carrier transgenic mice (C57BL/6 x C3H)^2^ and non-carrier mice (C57BL/6 x C3H) were both obtained from Jackson Laboratories and crossed. The F1 progeny of the cross were used for experiments. Lentivirus injections into ntg and PS19 mouse hippocampi were performed with stereotaxic surgery. Isoflurane anesthesia by inhalation was administered to the mice that were rested on a stereotax frame for surgery. To deliver lentivirus into bilateral hippocampi for each mouse the injection coordinates used from bregma were anterior-posterior: −2.0, medial-lateral ±1.5, and dorsal-ventral −1.8. A Hamilton syringe with needle was used for the injection of lentivirus into the hippocampus at 0.5 uL/min. A subcutaneous injection of buprenorphine was given to mice at the start of surgery, and two more buprenorphine injections were administered within 24 hrs after surgery for additional analgesic. Mice were housed in a pathogen-free barrier facility with a 12-hour light-dark cycle and provided with ad libitum access to water and food. Female and male mice were used in all experiments, and procedures followed the guidance and approval of the Institutional Animal Care and Use Committee (IACUC) of the Buck Institute.

### Electrophysiology

Whole-cell voltage clamp recordings were performed on human neurons in an external solution containing: 140 mM NaCl, 5 mM KCl, 10 mM glucose, 10 mM HEPES, 2 mM MgCl_2_, and 2.5 mM CaCl_2_, pH 7.4. Tetrodotoxin (0.5 μM, Abcam) was added to the external solution for recordings of miniature excitatory synaptic currents (mEPSCs). The patch pipette was filled with an internal solution containing: 120 mM CsCl, 10 mM HEPES, 1 mM EGTA, 5 mM NaCl, 1 mM MgCl_2_, 4 mM Mg-ATP, 0.3 mM Na_3_-GTP, pH 7.2 and the pipette resistance was 3-5 MΩ. Cells were voltage clamped at −60mV. The amplitude and frequency of mEPSCs were analyzed using SimplyFire^87^.

To make acute brain slices, mouse brains were dissected in cold sucrose solution containing (in mM): 210 sucrose, 2.5 KCl, 1.25 NaH_2_PO_4_, 25 NaHCO_3_, 7 glucose, 2 MgSO_4_, and 0.5 CaCl_2_ (perfused with 95% O2, 5% CO2 with ph ∼7.4). Horizontal 400 µm thick slices were cut using a vibratome (VT1000S, Leica) and placed into a recovery chamber for a 30 min incubation in artificial cerebrospinal fluid (ACSF) at 35°C containing (in mM): 119 NaCl, 2.5 KCl, 26.2 NaHCO_3_, 1 NaH_2_PO_4_, 11 Glucose, 1.3 MgSO_4_ (gassed with 95% O_2_, 5% CO_2_ ph ∼7.4). The recovery chamber with slices was then moved to room temperature with constant 95% O_2_, 5% CO_2_ bubbling maintained. Field potentials were recorded from the molecular layer of the dentate gyrus. A recording chamber held slices perfused with oxygenated ACSF solution kept at a constant 30°C. The recording electrode (3-4 MΩ resistance) was backfilled with ACSF and lowered 50 μm into the dorsal blade of the molecular layer of the dentate gyrus. The perforant pathway inputs to the dentate gyrus were stimulated with a concentric bipolar electrode (FHC) located 150 μm from the recording electrode. Input-output curves and maximal fEPSP slope were acquired with stimulus pulses elicited at an intensity range from 2.5 μA – 25 μA every 30 s with a 0.5 ms stimulus duration using a Model 2100 Isolated Pulse Stimulator (A-M Systems). We adjusted the stimulus intensity to 30% of the maximal fEPSP slope to record the 20 min baseline for LTP recordings in the presence of 100 μM picrotoxin (Sigma). After recording the baseline, the stimulus intensity was increased to 60% of the maximal fEPSP slope for the theta burst stimulation (TBS) which consisted of 10 theta bursts applied every 15 s and each theta burst contained 10 bursts (4 pulses, 100 Hz) every 200 ms. Following TBS, the stimulus intensity was adjusted back to the same magnitude as the baseline LTP recordings and field potentials were recorded for 60 min. Field EPSP slopes were normalized to the mean baseline LTP slope. Recordings were acquired with WinLTP software (version 1.11b, University of Bristol) using a Multiclamp 700B amplifier (Molecular Devices). Recordings and analyses were performed blind to genotype.

### Western Blot

Human brain tissue samples from temporal gyrus were homogenized in RIPA buffer containing 50 mM Tris, pH 7.5, 150 mM NaCl, 0.5% Nonidet P-40, 1 mM EDTA, 0.5% sodium deoxycholate, 0.1% SDS, 1 mM phenylmethyl sulfonyl fluoride, protease inhibitor cocktail (Sigma), phosphatase inhibitor cocktail (Sigma), 5 mM nicotinamide (Sigma) and 1 mM trichostatin A (Sigma). Tissue samples were homogenized with a dounce homogenizer, then sonicated 20 times with 1 sec pulses. The homogenates were centrifuged using a SW55 Ti rotor (Beckman Coulter) at 42,700 rpm for 15 min at 4°C. The resulting supernatant was used for western blot analysis as the RIPA-soluble fraction. A Bradford Assay (Bio-Rad) was used to measure the total protein concentration of each sample, and for each experiment, equivalent amounts of total protein across samples were loaded to run in a 4-12% gradient SDS-PAGE gel (Invitrogen). Proteins were transferred from the gel to a nitrocellulose membrane (GE Healthcare). Membranes were incubated in blocking solution containing 5% nonfat dry milk in TBST at room temperature for 1 hr which was followed by an overnight incubation with primary antibodies in TBST with 2% nonfat dry milk at 4°C, and three washes with TBST. Incubation with HRP-conjugated secondary antibodies was performed in TBST with 2% nonfat dry milk at room temperature for 1 hr, followed by three washes with TBST and labeling using an enhanced chemiluminescence (ECL) reagent (Pierce). The chemiluminescence from immunolabeling was exposed on film (Thermo Scientific), and images of the immunolabeling were quantified using Fiji software (ImageJ).

### Immunohistochemistry

Mice were anesthetized by an intraperitoneal injection of Avertin, and transcardial perfusion of ice-cold PBS was performed. Half brains were fixed in cold 4% PFA for 48 hours and following fixation, brains were washed and stored in PBS at 4°C. Before sectioning, brains were dehydrated in 30% sucrose with PBS for at least 48 hours then mounted on a Leica SM2010R microtome and cut into 30 μm thick sections. To begin the staining procedure, sections were washed three times for 10 min each in PBS and then permeabilized in 0.5% Triton X-100 in PBS for 10 min. Sections were incubated in blocking solution containing 10% normal goat serum in PBS-T (0.5% Triton X-100) for 1 hour at room temperature. Sections were incubated in primary antibodies mixed in PBS-T with 3% normal goat overnight at 4°C. After three PBS-T washes for 10 min each, secondary antibodies were applied in PBS-T with 3% normal goat serum for 1 hour at room temperature. After three washes in PBS-T for 10 min each, sections were mounted on glass slides with Prolong Gold. Antigen retrieval was performed for eIF4B immunolabeling in brain sections. Sections were first washed three times for 5 min each in PBS, then incubated in 10 mM sodium citrate pH 6.0 for 3 min at 95°C. Sections were allowed to cool for 20 min at room temperature then washed another three times for 5 min each in PBS. After washes, sections were incubated in blocking solution and the remainder of the standard staining protocol was followed as described above.

### Proximity Ligation Assay (PLA)

The manufacturer’s protocol for the NaveniFlex MR kit was followed to perform PLA on fixed HEK293 cells or human iPSC-derived neurons grown on glass coverslips. Coverslips were blocked for 1 hour at 37°C then incubated in primary antibody for 1 hour at 37°C. After three washes in TBS-T (0.15% Tween), coverslips were incubated with Navenibody secondary antibodies for 1 hour at 37°C. After secondary antibody labeling, reactions A, B, and C were completed as described in the manufacturer’s protocol. Coverslips were mounted on glass slides. We imaged PLA signal at regions with similar cell densities across experimental groups in human iPSC-derived neurons and quantified the density of PLA puncta in the entire acquisition window. In HEK293 cells, the PLA signal was imaged in GFP-positive cells. The PLA in HEK293 cells was not as punctate as the PLA in human neurons due to the higher concentration of PLA signals in the HEK293 cells. Consequently, the mean intensity of PLA signal, rather than the density of puncta, was quantified in HEK293 cells. Image acquisition and analysis were performed blind to the treatment and genotype.

### Polysome Profiling

Human iPSC-derived neurons were plated at a density of 8 ×10^6^ cells on 10 cm plates co-plated with 500,000 rat astrocytes. Polysomes were acquired from human neurons at 7-8 weeks of maturation. Human neurons were induced with cLTP and 45 min post induction, cycloheximide (1 µg/mL final concentration) was added to cultures and incubated for 10 min at 37°C. After cycloheximide treatment, neurons were washed 2X in ice-cold PBS. Neurons were homogenized in lysis buffer (10 mM MgCl_2_, 100 mM NaCl, 20 mM Tris, pH 7.5, 0.4% NP-40, 1 ug/mL cycloheximide) supplemented with Complete EDTA-free protease inhibitor. Homogenate was centrifuged for 10 min at 4°C at 15000 rpm. Supernatant was pipetted onto an ice-cold 10-50% sucrose gradient (10 mM MgCl_2_, 100 mM NaCl, 20 mM Tris, pH 7.5) and centrifuged in a SW41Ti rotor at 38000 rpm for 2 hrs at 4°C. Fractions were collected with a Biocomp Gradient Fractionator and profiles were generated by measuring the optical density at 254 nm light with a Biorad Econo UV monitor. The area under the curve was calculated from the optical density measurements made on the polysome fractions.

### RNA Sequencing Analyses

RNA from polysome fractions were extracted using TRIzol LS reagent (Invitrogen) and stored at −80°C. Equal amounts of RNA from polysome fractions 16, 17, and 18 were pooled for RNAseq from each sample. Sequencing was performed at the Genomics Core Facility at the Icahn School of Medicine at Mount Sinai. RNA samples were DNase treated and then validated for quality with Qubit RNA quantification and Agilent BioAnalyzer for RNA integrity (RIN > 8). Sample libraries were prepared using 5 ng of total RNA from each sample with a Takara-seq v4 Ultra Low Input kit and sequenced on an Illumina NovaSeq 6000 sequencing system. We sequenced 50 million reads per sample to investigate differential gene expression between tauWT and tauV337M_iso_ neurons both treated with cLTP.

Fastq reads were trimmed using fastq_quality_trimmer (-Q 33, -t 20, -l 20) and filtered using fastq_quality_filter (-Q 33, -q 20 -p 100) from the FastX toolkit^88^. Remaining read pairs were aligned to the GrCH38 assembly of the human genome using hisat2^89^. Output sam files were converted to bam and sorted using samtools^90^ and read counts were calculated using featurecounts^91^. PCA plots were made using the DESeq2^92^ package and including the top 200 variable genes. Gene Ensemble ids were converted to gene symbols using the org.Hs.eg.db ^93^ database and the mapIds function in the AnnotationDbi^94^ package. Expression differences between mutant and wildtype were calculated using edgeR^95^. Rank-rank hypergeometric overlap analysis was completed using the RRHO2^96^ package using genes ranked by log fold-change. Gene Ontology (GO) term enrichment analyses for Cellular Compartments were performed using ClueGO version 2.5.4 in Cytoscape version 3.7.1^97,98^. ClueGO statistics revealed enriched networks with p-values < 0.05 from a two-sided hypergeometric test with Bonferroni correction. Synaptic Gene Ontologies (SynGO) version 1.2 was used to evaluate the transcripts of synaptic proteins that were identified^99^.

To perform RT-PCR on the same samples that were used for RNA-seq, cDNA was generated from the RNA extracted from polysome fractions #16, 17, 18 using the iScript cDNA Synthesis Kit (BioRad). The following PCR primers were used for RT-PCR: PKMζ Forward: CTT ACA TTT CCT CAT CCC GGA AG; PKMζ Reverse: TTC ACC ACT TTC ATG GCG TAA A; Shank3 Forward: GGG ATC ACC GAC GAG AAT GG; Shank3 Reverse: TGT CTG CCC CAT AGA ACA GC; Actin Forward: ACA CCC TTT CTT GAC AAA ACC T; Actin Reverse: CGC ATC TCA TAT TTG GAA TGA CT. Agarose gel electrophoresis was used to analyze PCR products. After subtracting background, gel band intensities were quantified using ImageJ software (NIH).

### Mouse Behavior

For the object-context discrimination test, mice were habituated for 30 min in the behavior room before the start of testing. During the sample phase, mice either explored Context 1 which contained two identical objects (X1, X2) in a white box that was cleaned with 70% ethanol or Context 2 which contained a different set of objects (Y1, Y2) in a white box with black and white checkered wallpaper that was cleaned with 1% acetic acid. Each mouse had two (one for each context) 10 min sessions that were 30 min apart. After the second sample phase session, mice rested in their home cages for 4 h. Following the rest period, one of the Y objects in Context 2 was replaced with an incongruent X object, and one of the X objects in Context 1 was replaced with an incongruent Y object. The mice were split equally based on genotype and sex, so half the mice were placed in Context 1 and the other half in Context 2 for the test phase. We hand-scored the time mice spent exploring the congruent and incongruent objects in a 10 min session for each mouse. Videos of the sessions were manually scored by an experimenter who was blind to mouse genotype and treatment.

For the Morris water maze (MWM) test, visual cues were placed around a pool with a diameter of 1.2 m and filled with water maintained at a temperature of 22°C containing white tempera paint. On day one, mice were trained with 4 trials to find a hidden square platform (14 cm x 14 cm) in the middle of a rectangular channel at 1.5 cm below the water surface. The next day mice started hidden platform training where each mouse had 1 min to find and sit independently on the platform for 10 s. Mice performed 2 trials per day for 5 days of hidden platform training. The next day (24 h) after hidden platform training, the platform was removed from the pool and mice were tested in a single probe trial for 60 s. Each mouse was recorded and tracked using EthoVision XT software (Noldus). The experimenter was blind to the genotype and treatment of mice. All behavior experiments were performed during daylight hours.

### Human samples

Samples of human brain tissue were obtained from the Neurodegenerative Disease Brain Bank at the University of California San Francisco. Histological and immunohistochemical methods were performed as described^100,101^ for neuropathological diagnoses that were made following consensus diagnostic criteria^102,103^. Clinical and neuropathological diagnoses were used to select the cases. Middle temporal gyrus was dissected from frozen brain tissue from control cases (n=7; ADNC ≤ A1B1C1; mean age: 85.71 ± 6.05 years; sex: 5F, 2M) and AD cases (n = 7; ADNC A3B3C3; mean age: 85.71 ± 2.87 years; sex: 1F, 6M).

### Statistical Analyses

Student’s *t*-test was applied to evaluate differences between the means of two groups. A one-way or two-way ANOVA with Bonferroni multiple comparison *post*-hoc analyses were used to test the differences between means of multiple groups. The fEPSP slope and fiber volley amplitude analyses across stimulus intensities in dentate gyrus and the analysis of spatial learning during the hidden platform training of the MWM were both tested by repeated measures two-way ANOVA with Bonferroni multiple comparison.

