## Supplementary figures and images for "Pathogenic tau inhibits synaptic plasticity by blocking eIF4B-mediated local protein synthesis"

### Supplementary figures

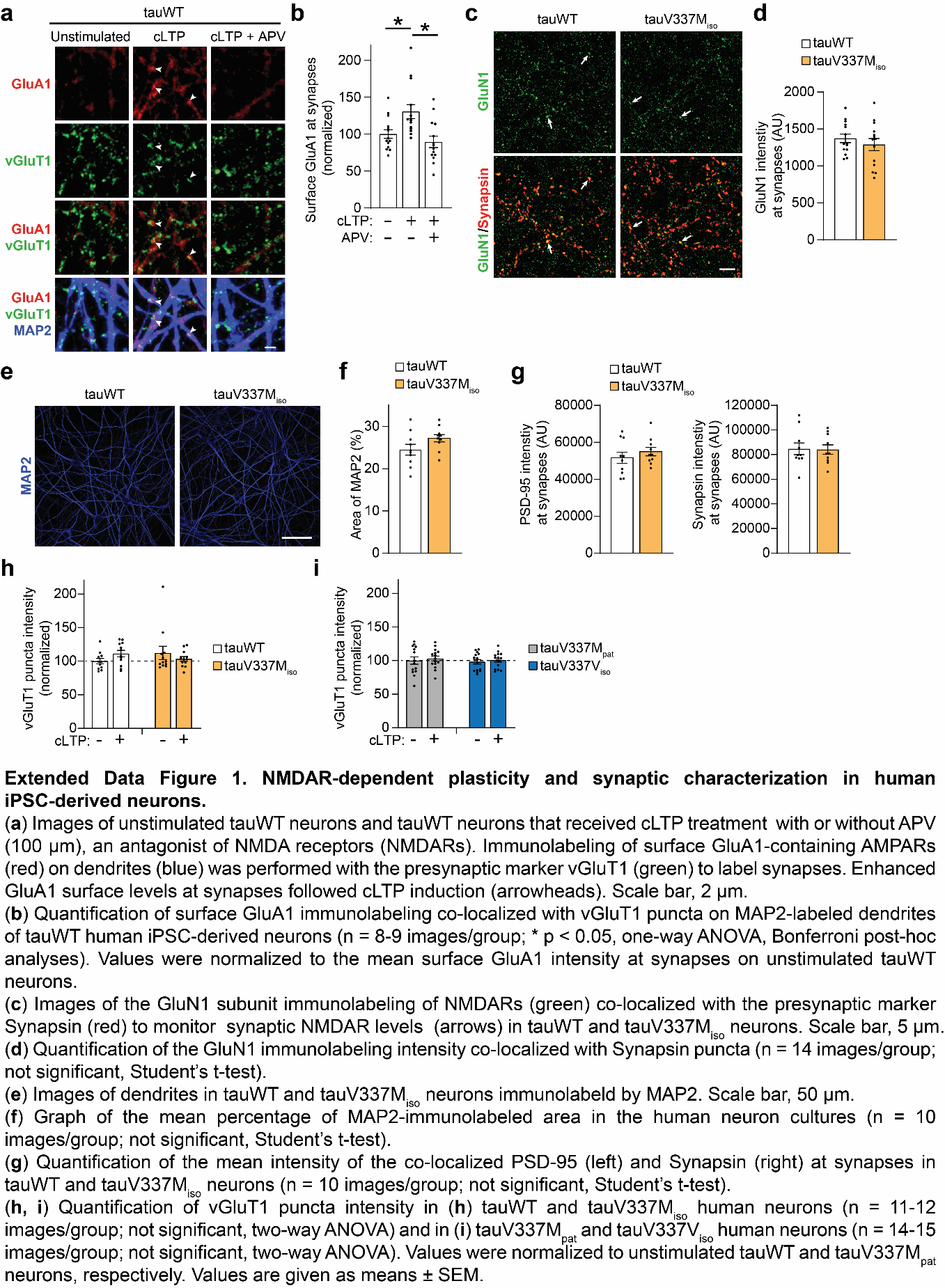

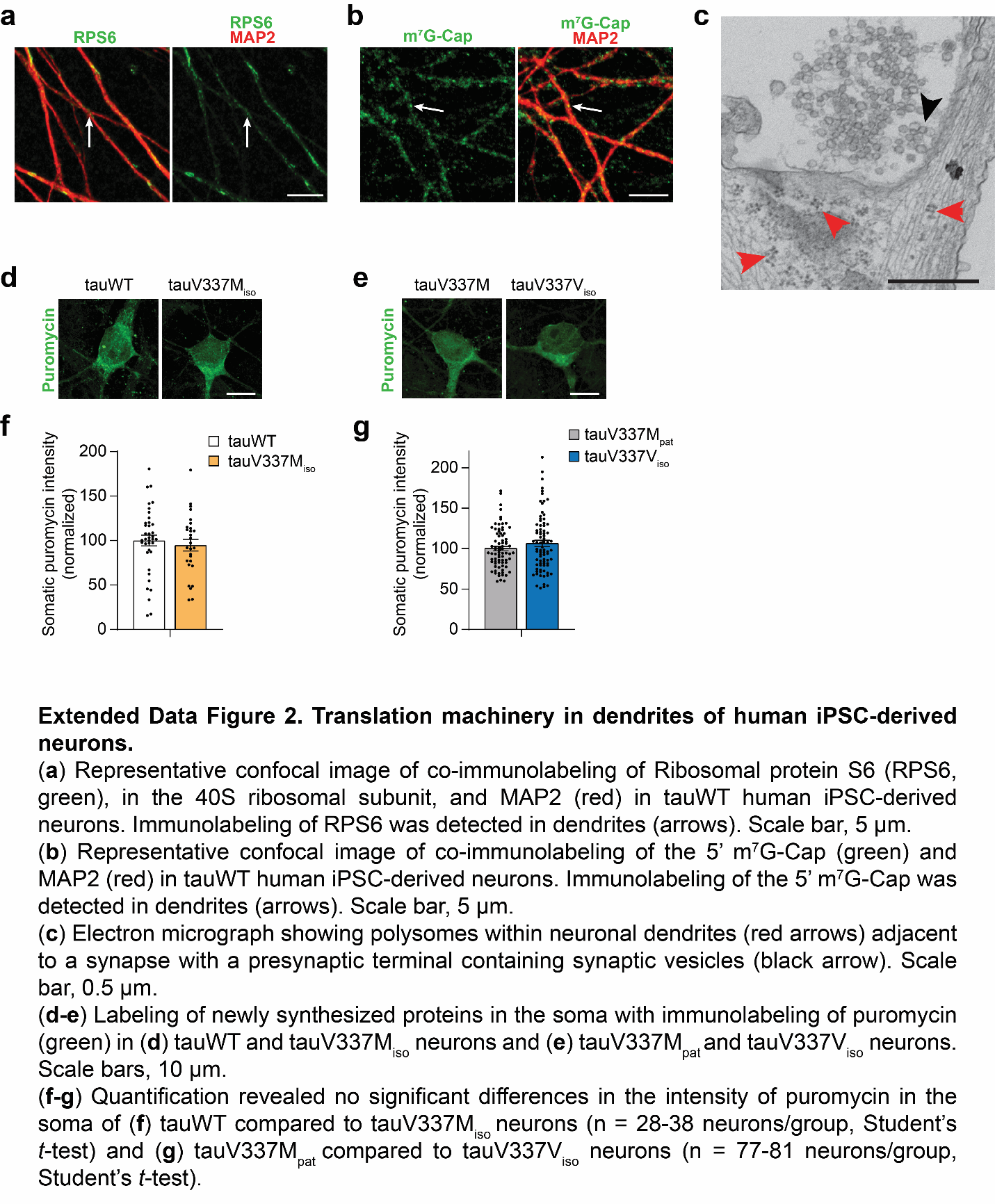

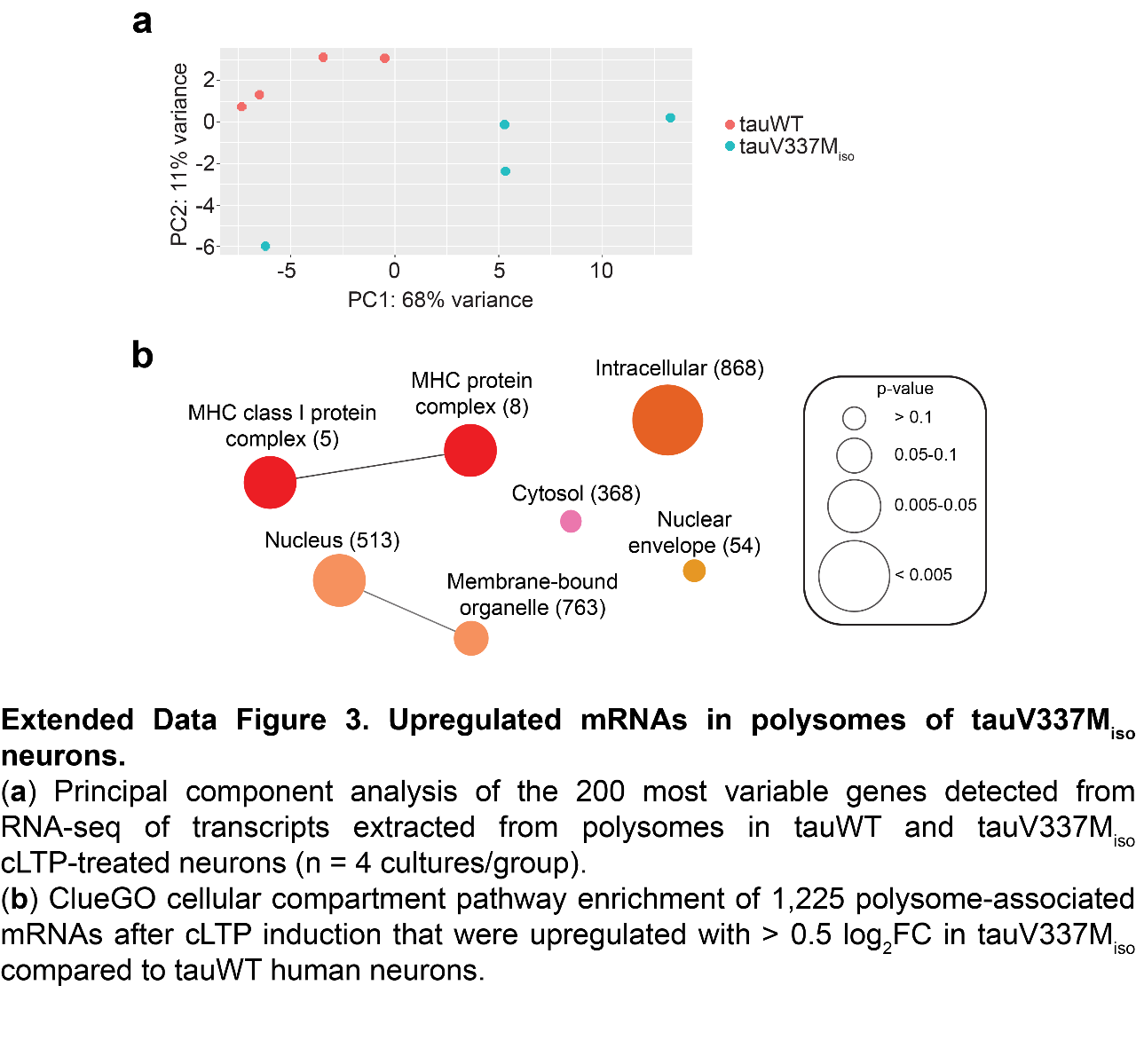

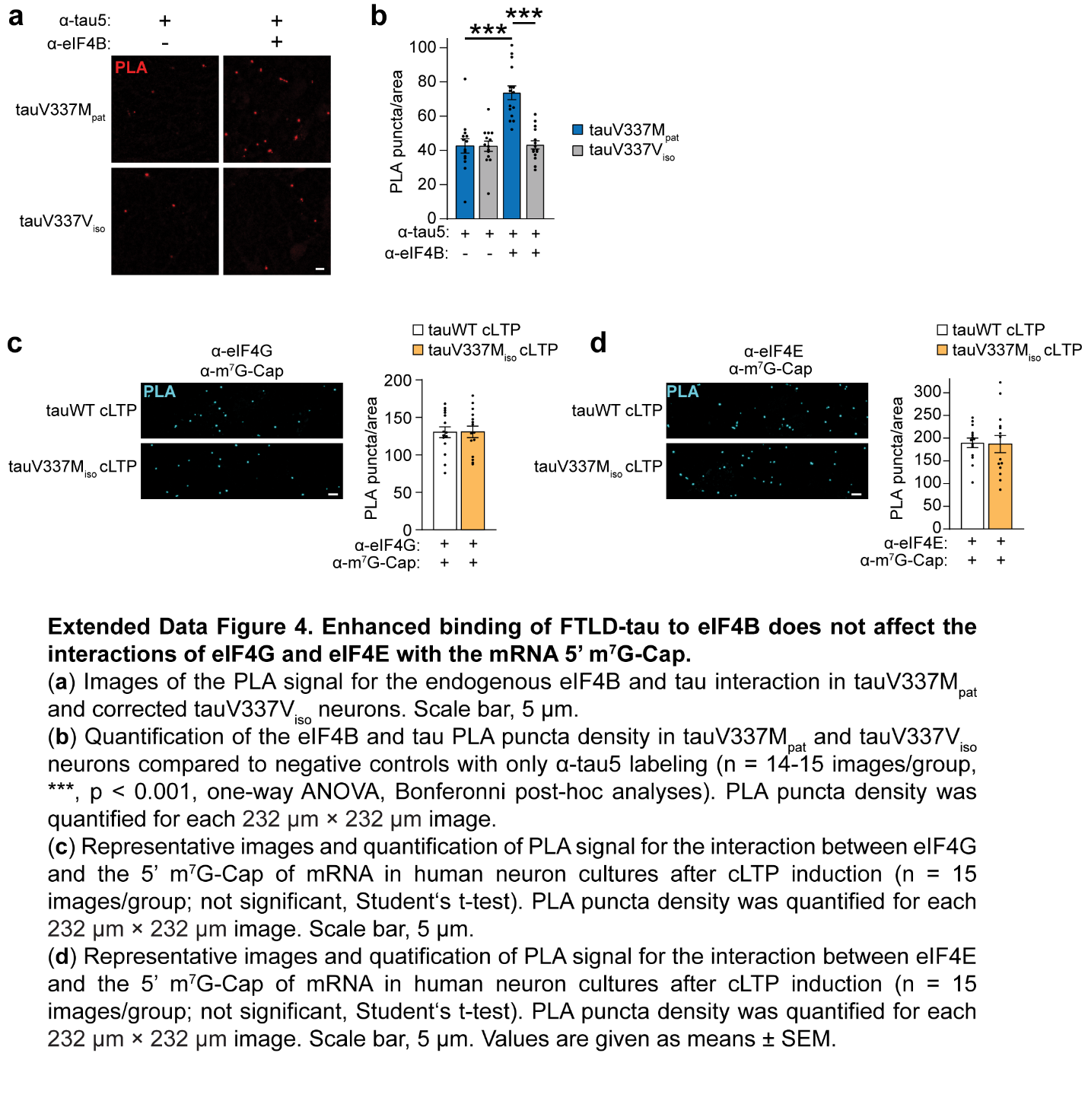

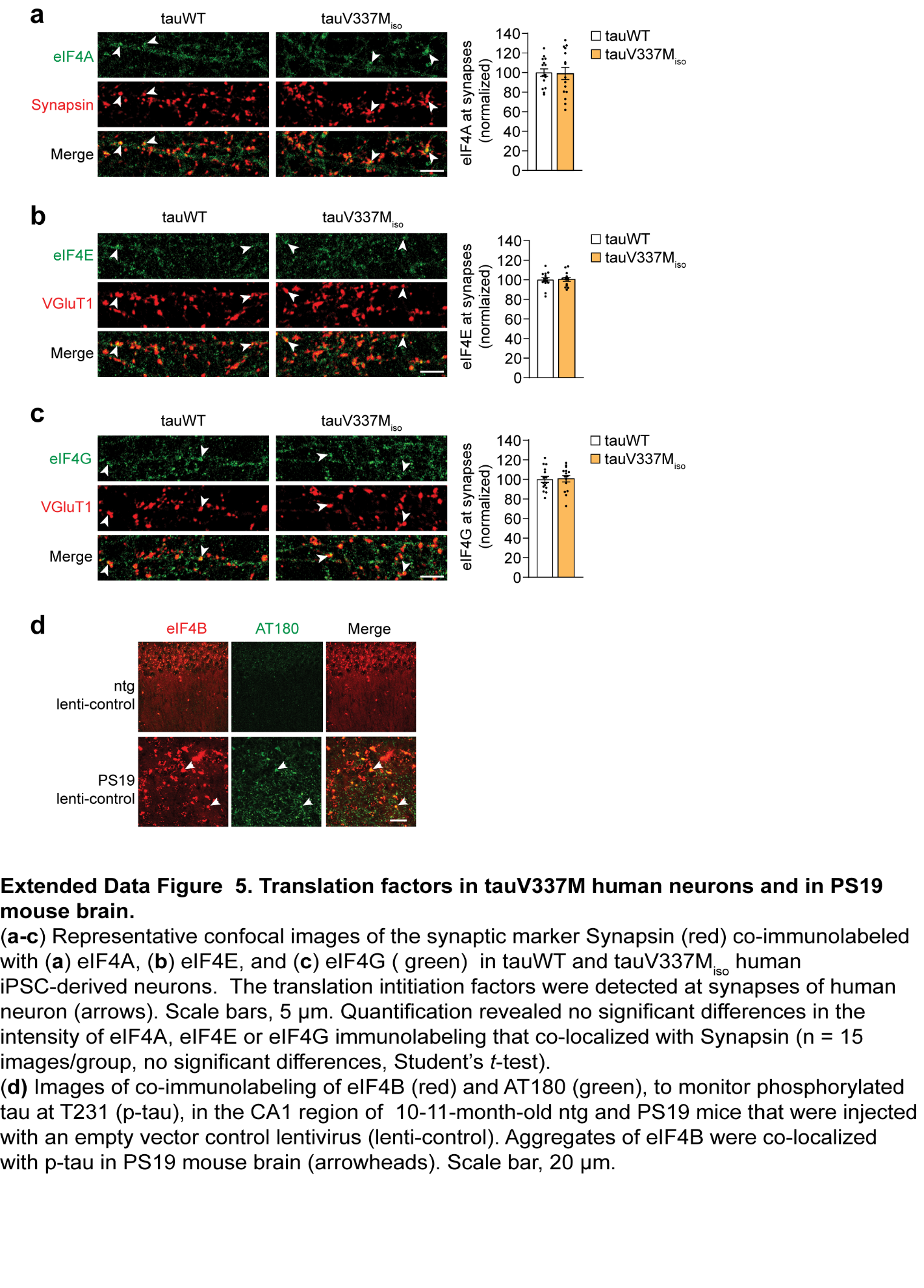

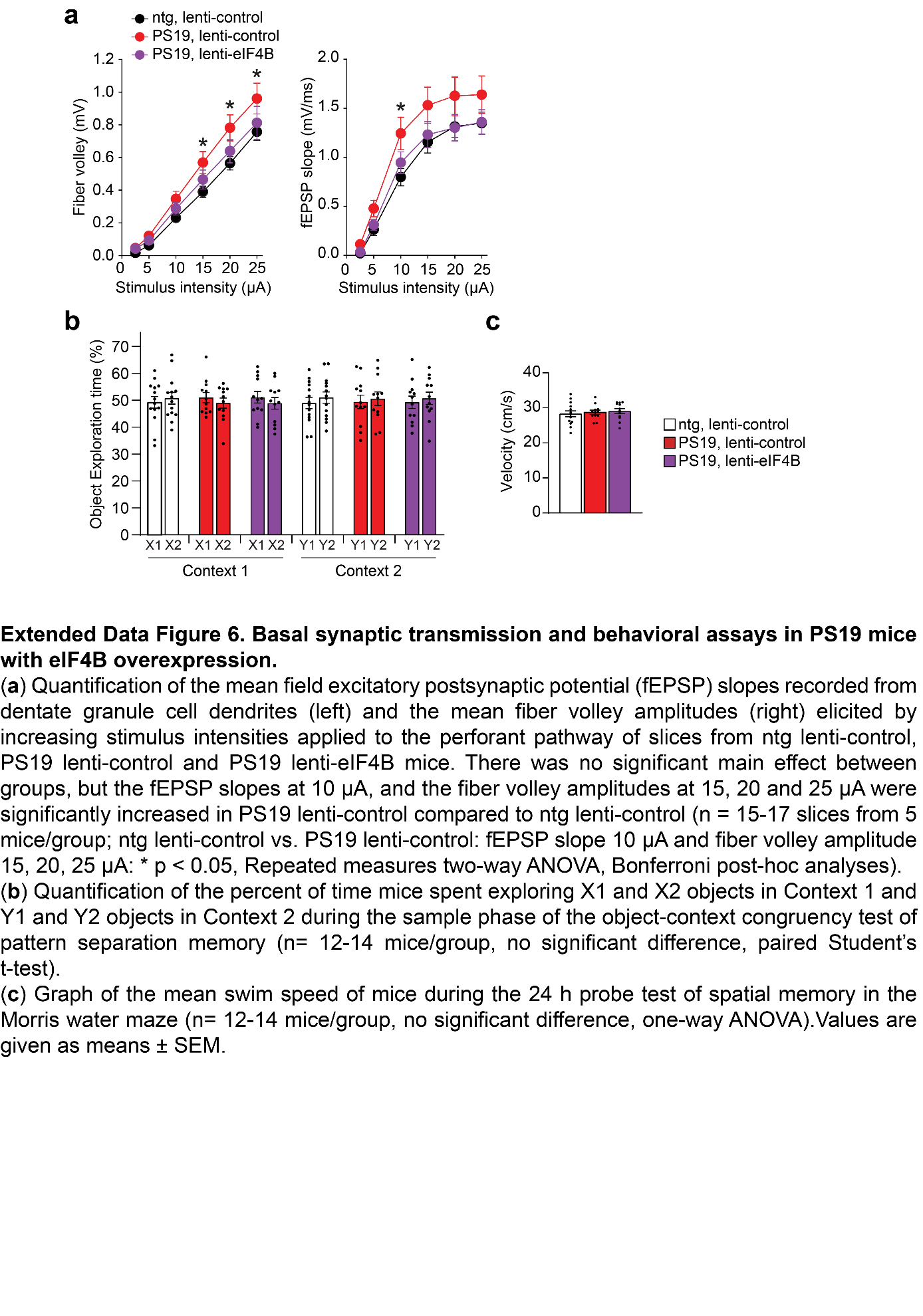

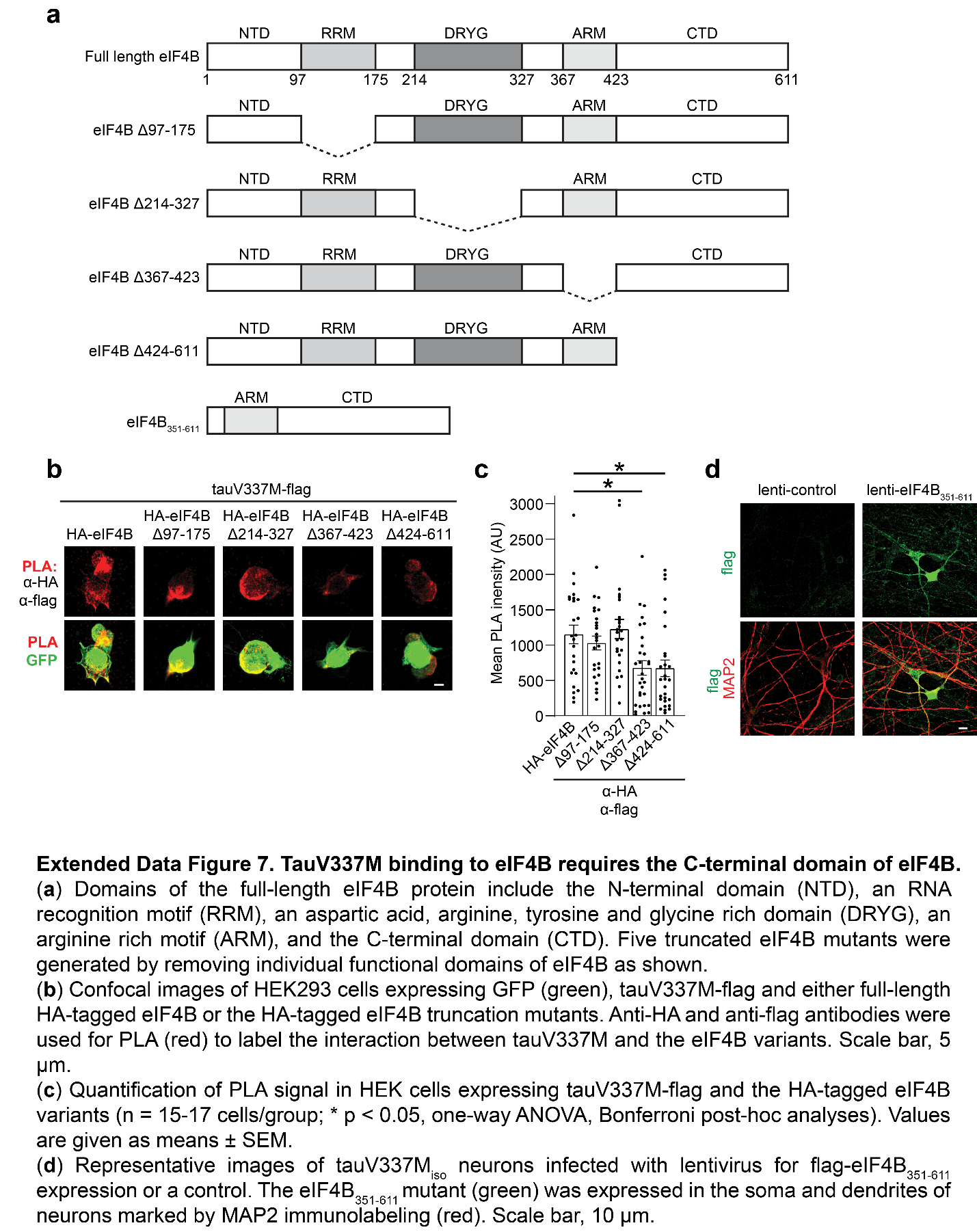
